# The projected economic and health burden of uncontrolled asthma in the United States

**DOI:** 10.1101/516740

**Authors:** Mohsen Yaghoubi, Amin Adibi, Abdollah Safari, J Mark FitzGerald, Mohsen Sadatsafavi, *for the Canadian Respiratory Research Network*

## Abstract

**Rationale:** Despite effective treatments, a large proportion of asthma patients do not achieve sustained asthma control. The ‘preventable’ burden associated with lack of proper control is likely taking a high toll at the population level.

**Objective:** We predicted the future health and economic burden of uncontrolled asthma among American adults for the next 20 years.

**Methods:** We built a probabilistic model that linked state-specific estimates of population growth, asthma prevalence rates, and distribution of asthma control levels. We conducted several meta-analyses to estimate the adjusted differences in healthcare resource use, quality-adjusted life years (QALYs), and productivity loss across control levels. We projected, nationally and at the state-level, total direct and indirect costs (in 2018 USD) and QALYs lost due to uncontrolled asthma from 2019 to 2038 in the United States.

**Measurements and Main Results:** Over the next 20 years, the total undiscounted direct costs associated with suboptimal asthma control will be $300.6 billion (95% confidence interval [CI] $190.1 – $411.1). When indirect costs are added, total economic burden will be $963.5 billion (95%CI $664.1 – $1,262.9). American adolescents and adults will lose 15.46 million (95%CI 12.77 million – 18.14 million) QALYs over this period due to suboptimal control of asthma. In state-level analysis, the average 20-year per-capita costs due to uncontrolled asthma ranged from $2,209 (Arkansas) to $6,132 (Connecticut).

**Conclusion:** The burden of uncontrolled asthma will continue to grow for the next twenty years. Strategies towards better management of asthma may be associated with substantial return on investment.

## Introduction

Asthma is a very common chronic disease globally. The prevalence of asthma has increased over the last decade in many regions of the world(1). Despite the fact that asthma imposes a substantial burden on patients and healthcare systems, it has not yet been identified as a healthcare priority in many countries(1). In the U.S., there are approximately 26 million patients with physician-diagnosed asthma(2). Asthma cost the U.S. economy an estimated $81.9 billion USD in 2013 alone(3).

Conventional wisdom suggests that asthma is not a curable disease. Indeed, evidence indicates that airway hyper-responsiveness and inflammation persist in individuals whose asthma has been dormant for many years(4). Therefore, the contemporary asthma management paradigm is based on achieving symptom control and reducing the risk of exacerbations(5). It is widely accepted that through avoidance of triggers and use of inhaled anti-inflammatory agents (namely inhaled corticosteroids [ICS]), asthma can be controlled and exacerbation risk can be significantly reduced in the majority of patients(6). Achieving asthma control is associated with improvement in quality of life, reduction in medical costs, and better work performance(7,8). Unfortunately, the reality of asthma care is highlighted by poor adherence to treatments and other disease management modalities (e.g., avoidance of triggers), resulting in a significant proportion of asthma patients experiencing suboptimal asthma control(9).

A good understanding of the future burden of diseases can support the search for efficiency and equity in healthcare. Many studies have estimated the total burden of asthma in the U.S.(1,3,10). However, given the focus of contemporary asthma management is on achieving asthma control, the relevant figure of merit for policymaking and prioritizing future research is the burden due to suboptimal asthma control, rather than the burden of asthma itself. In a small minority of patients, achieving symptom control can be difficult or out of reach (11). However, given the availability and efficacy of inexpensive management strategies to control asthma in the majority of patients, the excess burden between uncontrolled and controlled asthma is largely ‘preventable’. Such burden can be considered as the maximum space available for contemporary asthma management strategies to reduce asthma burden at the population level. The resources required for interventions and programs aimed at improving asthma control can be juxtaposed against their estimated impact on the burden of suboptimal asthma control to evaluate whether such programs are worth implementing.

The purpose of the present study was to document the current economic and health burden as well as project the future preventable economic and health burden associated with suboptimal asthma control among the U.S. adolescent and adult population for the next 20 years. We answered the question “*how much costs could be saved, and quality of life could be improved, if all adolescent and adult asthma patients in the U.S. achieve symptom control in the next 20 years?”*

## Methods

To enable projections, we reconciled evidence from multiple sources into a time-in-state computer model of asthma. The projection period was from 2019 to 2038 (20 years). We adopted a societal perspective in the primary analysis; thus, costs were included no matter who had incurred them. The analyses were performed for the entire U.S. population ≥14 years of age, as well as at the state level. We defined asthma control according to the score on the Asthma Control Test (ACT)(12). ACT was used because major sources of evidence for this study used this instrument(13–19). This test classifies the patient’s asthma status into poorly controlled (scores ≤ 15; 29.3%), not well controlled (score 16-19; 25.1%), and well controlled (score 20-25; 45.7%). As the focus was on achieving (well) controlled asthma, the outcomes of very poorly controlled and not well controlled were combined into one category (henceforth referred to as ‘uncontrolled’). The supplementary material provides details of the methodology. *Table 1* provides point estimates and probability distribution assigned to each model parameters.

**Table 1.** Model input parameters

| Parameters | Point estimate | Probability distribution† | Source |
| --- | --- | --- | --- |
| Forecast of U.S. population growth |  |  | US national population projection (20,21) (see supplementary material) |
| Prevalence of asthma in U.S. across age ,sex and state |  |  | Global Burden of Diseases(22,23) (see supplementary material) |
| Association between asthma control and age/sex‡<br>Well-controlled (vs. very poorly controlled )<br>age<br>sex<br>Not well-controlled (vs. very poorly controlled )<br>age<br>sex | 1.09<br>0.74<br>0.99<br>1.10 | Log Normal†(0.08,1.23)<br>Log Normal(-0.30,1.32)<br>Log Normal(-0.01, 1.28)<br>Log Normal(0.09, 1.47) | Based on model calibration (see supplementary material) |
| Pooled odd ratio of health care provider Uncontrolled (vs. well-controlled) | 1.86 | Log Normal(0.62,0.53)† | Meta-Analysis (see supplementary material) |
| Pooled odd ratio of emergency visit Uncontrolled (vs. well-controlled) | 1.44 | Log Normal(0.36,0.05) | Meta-Analysis |
| Pooled odd ratio of hospitalization Uncontrolled (vs. well-controlled) | 1.54 | Log Normal(0.43,0.29) | Meta-Analysis |
| Pooled odd ratio of medication use Uncontrolled (vs. well- controlled) | 1.58 | Log Normal(0.45,0.32) | Meta-Analysis |
| Excess direct medical costs ( per person-2018US\$)* Uncontrolled (vs. well-controlled) | 1,349 | Normal† (1,349, 480) | Estimated through calibration, combining distribution of asthma controls and resource use ratios (above rows) with estimate of overall costs of asthma (3) |
| Pooled mean difference of overall work impairment (%) between uncontrolled and well-controlled | 12.70% | Log Normal(2.5,3.3) | Meta-Analysis |
| Excess Indirect medical Costs (per person-year, 2018US\$) between uncontrolled and well-controlled | 3,350 | Normal(3,350,886) | Estimated through calibration, combining distribution of asthma controls and overall work impairment (above row) with estimate of average income (24) |
| Pooled mean difference of QALY lost between uncontrolled and. controlled | 0.07 | Log Normal(-2.65,0.01) | Meta-Analysis |

\*All costs are in USD 2018.
†Normal(x, y): Normal distribution with mean x, and standard deviation y, lognormal(x, y), log normal distribution with mean x and standard deviation y for the log-transformed values
‡We modeled probability of three levels of control to align with calibration process and pooled “very poorly control” and “not-well control” into “uncontrolled group” in order to report outcome.

### Data sources

We used the following six major sources of evidence to populate the model. Details of the estimated parameters and methodologies are provided in Supplementary Materials, Section 1.

1. Forecasts of population growth and aging during the projection period, nationally and for each state, were derived from the National Population Projections conducted by the Census Bureau - Population Division(20,21).We estimated the midpoint of two sets of national population projections based on the 2010 Census for bases case estimate.
2. Estimates of the prevalence of asthma, stratified by age and sex for each state, were obtained from the Global Burden of Disease (GBD) studies in 2016(22,23).These studies used the systematic analysis of published literature and other sources to estimate, using a consistent methodology, the burden of several health conditions including asthma(25).
3. Distribution of control levels in the asthma population, stratified by sex and age groups, were derived using calibration techniques from a recent study based on the U.S. National Health and Wellness Survey (NHWS) between 2011 and 2013(13). We estimated the sex-and age-specific distribution of control levels by solving the coefficients of a multinomial logit equation (with prevalence of asthma control levels as the outcome, and sex and age as independent variables) such that the prevalence of asthma control levels matches NHWS estimates (*Table 1*). Details of calibration techniques are provided in the Supplementary Materials (Section 1.3).
4. Healthcare resource use and quality-adjusted life years (QALY) differences across control levels were based on dedicated literature reviews and meta-analyses. We retrieved all relevant U.S.-based studies that assessed the adjusted association between asthma control and healthcare resource utilization, overall work impairment (indirect costs), and health-related quality of life. Studies were included if they controlled for potential confounding variables. Random-effects models were used to estimate the pooled adjusted odd ratios (OR) and 95% confidence interval (95%CI) of the association between asthma control and rate of healthcare provider visits, emergency visits, hospitalizations, and medication use. In addition, we used random-effects models to estimate adjusted pooled mean difference of overall work impairment in terms of the percentage of work hours lost due to sub-optimal asthma control. Finally, we pooled mean differences in QALYs between the uncontrolled and controlled groups. Details of the search strategy and meta-analysis are provided in the Supplementary Materials (Section 1.4).
5. We performed model calibration to convert estimates of resource use to direct costs. This method combines the distribution of asthma control levels in the population, ratio of resource use between uncontrolled and controlled asthma, and total direct costs of asthma in the U.S., to solve for the costs of uncontrolled versus controlled asthma. The first two components were obtained as described above. For total costs of asthma in the U.S., we relied on a recent large study (n=214,000) by the Centers for Disease Control and Prevention(3). This study used population-based sampling to estimate the overall costs of asthma. We solved for the costs of uncontrolled and controlled asthma that produced the desired ratio between the two that matched the results of our meta-analysis of resource use, and summed up to the total costs of asthma. Details of this methodology are provided in the Supplementary Materials (Section 1.5).
6. To estimate indirect costs suboptimal asthma control, we obtained the monetary value of productivity loss based on age-and sex-specific wages as reported by the Bureau of Labour Statistics(24). These estimates were combined with the results of the meta-analysis of productivity loss differences across control levels.

### Analysis

All projections are made for the period of 2019 to 2038. The primary projections are made for the entire U.S. adolescent and adult asthma population. State-level projections are provided as secondary results. In the main analysis, we estimated undiscounted total direct costs, indirect costs, and QALYs lost attributable to uncontrolled asthma. In a sensitivity analysis we calculated outcomes after applying a 3% annual discount rate, as recommended by panel on Cost-Effectiveness in Health and Medicine(26). All costs were adjusted to 2018 U.S. dollars using historical inflation rates(27). Uncertainty was modeled by assigning probability distributions to all input parameters (e.g., based on the reported 95%CI) and was propagated to the uncertainty in the projections using Monte Carlo simulation: within each simulation loop, we randomly drew from the distribution of all model parameters, performed model calibrations as described above, and calculated the outcomes. Uncertainty was presented in terms of 95%CIs around point estimates of projections.

## Results

### Systematic review and meta-analysis of resource use across control levels

We identified 10 studies that reported on the adjusted differences in direct costs, indirect costs, or QALYs across control levels in the U.S. Forest plots for adjusted odds ratios (ORs) and mean differences are shown in *Figure 1* and *Figure 2*, respectively. For healthcare resource use, the pooled adjusted ORs in the uncontrolled versus the controlled group were as follows: 1.86 (95%CI 1.34 – 2.38; 5 studies) for physician visits, 1.44 (95%CI 1.39 – 1.49; 5 studies) for emergency room visits, 1.54 (95%CI 1.25 – 1.80; 3 studies) for asthma-related admissions, and 1.58 (95%CI 1.26 – 1.90; 3 studies) for medication use (all P<0.001). Combining these ratios with the distribution of control levels and overall costs of asthma, the estimated excess direct costs associated with uncontrolled versus controlled asthma were $1,349 (95%CI $868 – $1,829) per patient-year. The pooled standardized mean difference of overall work impairment between the uncontrolled and controlled groups was 12.7% (95%CI 9.4% – 16.0%; P<0.001; 3 studies). Assuming 52 workweeks in a year, this translates to a loss of 6.6 extra weeks of productivity loss per year for each patient with uncontrolled asthma. Excess indirect costs between the two groups were estimated to be $3,350 (95%CI $2,464 – $4,236) per patient-year.

**Figure 1.**
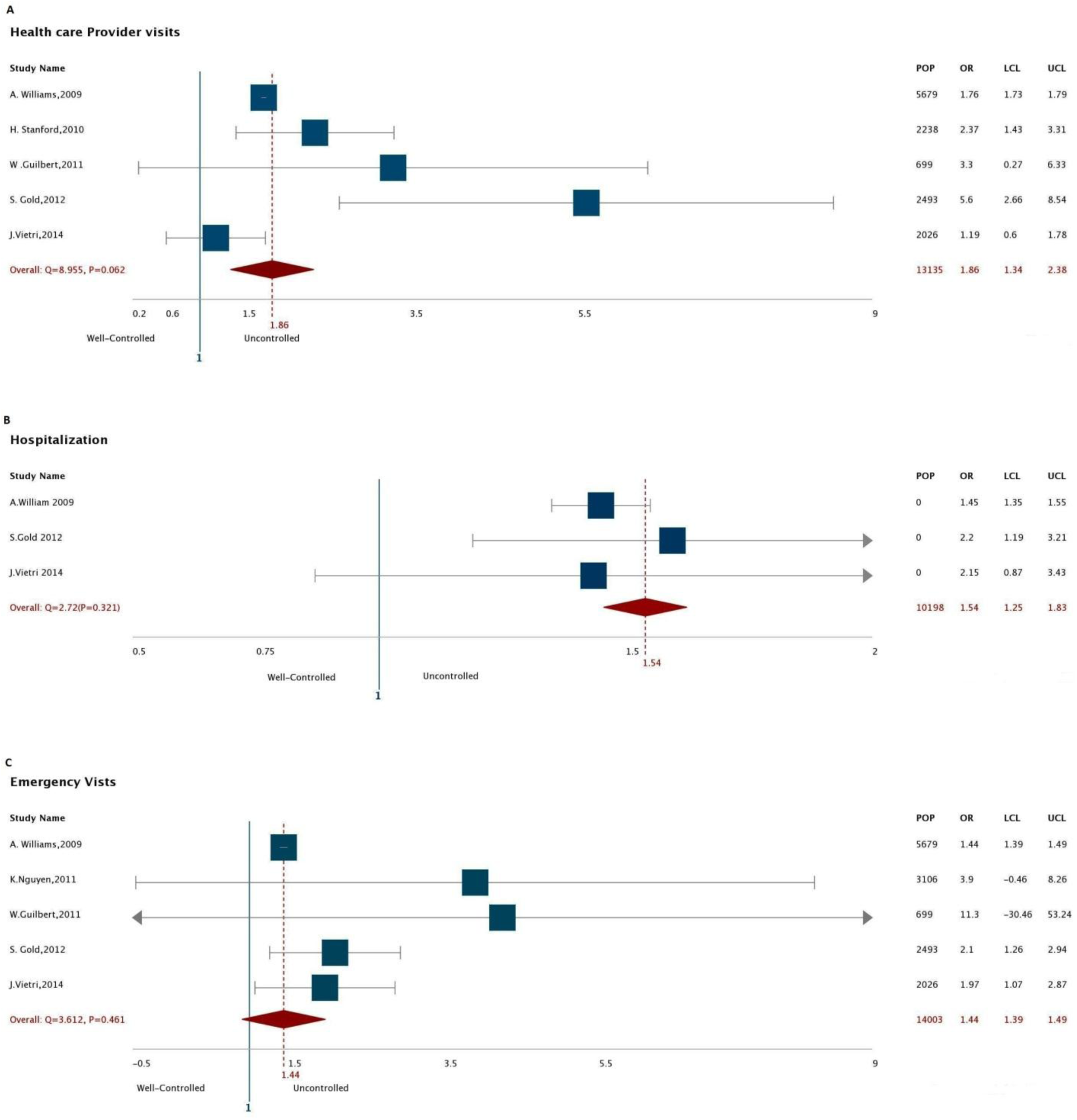
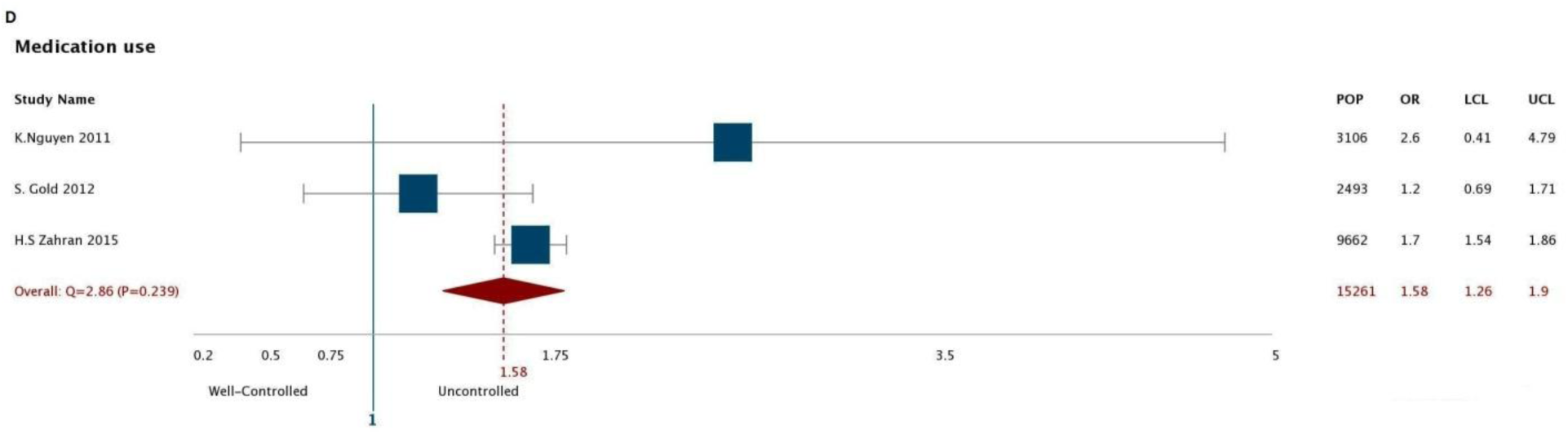
Forest plots of odds ratio of health care provider visit (A), Hospitalization (B), Emergency visit (C) and Medication use (D) across level of controls. Pop=Population; OR= Odd Ratio; CL= Lower Confidence Level; UCL= Upper Confidence Level.

**Figure 2.**
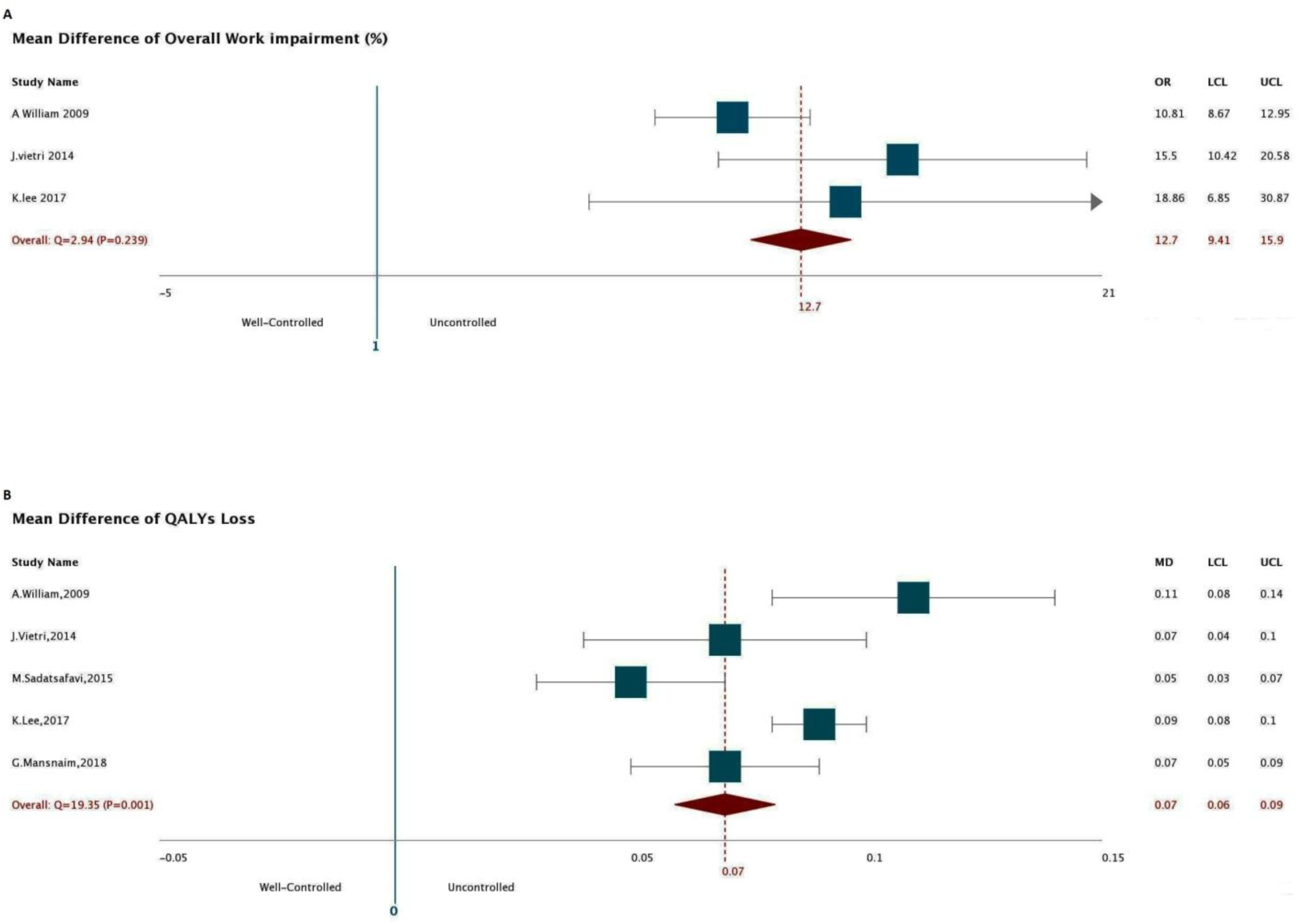
Forest plots of adjusted mean difference of overall work impairment (A) and QALYS loss (B) across level of controls. Pop=Population; OR= Odd Ratio; CL= Lower Confidence Level; UCL= Upper Confidence Level

Finally, the pooled estimate of the mean reduction in QALY values was 0.07 (95%CI 0.06 – 0.09; P<0.001; 5 studies). Further details on these results are provided in the Supplementary Materials (Table E4-E7 and FigureE1).

### Projection of burden of suboptimal asthma control

The size of the U.S. adolescent and adult population was projected to increase by 11%, from 266.87 million in 2019 to 298.22 million in 2038. In 2019, there will be 15.88 million adolescent/adult asthma patients in the country, which is expected to increase to 17.65 million by 2038, representing 10% growth; 62% of asthma patients will be women. During the 20-year projection window, in 175.32 million patient-years (52% of total patients-years of asthma), asthma will be sub-optimally controlled. Total undiscounted direct costs of asthma across all control levels combined over 20 years will be $1,537 billion. The total undiscounted 20-years direct costs, indirect costs, and QALYs lost associated with sub optimal controlled of asthma within age and sex group are provided in *Table 2*.

**Table 2.** The undiscounted projected 20-years direct costs, indirect costs and QALYs lost associated with suboptimal control of asthma within age and sex group

| Sex | Age | Excess Direct Costs[SD] (Million \$) | Excess Indirect Costs[SD] (Million \$) | Excess QALYs lost[SD] |
| --- | --- | --- | --- | --- |
| Female | 15 to 19 | 14,236[5,254.9] | 35,138[10,062.6] | 732,193[128,686] |
| Female | 20 to 24 | 14,744[5,434.2] | 36,392[10,395.7] | 758,322[132,503] |
| Female | 25 to 29 | 14,965[5,507.7] | 36,935[10,526.5] | 769,649[133,759] |
| Female | 30 to 34 | 14,107[5,185.7] | 34,817[9,902.6] | 725,533[125,499] |
| Female | 35 to 39 | 13,295[4,882.1] | 32,811[9,316.0] | 683,734[117,820] |
| Female | 40 to 44 | 12,431[4,561.8] | 30,679[8,699.6] | 639,309[109,879] |
| Female | 45 to 49 | 12,976[4,760.0] | 32,022[9,074.2] | 667,315[114,566] |
| Female | 50 to 54 | 13,347[4,896.7] | 32,938[9,333.3] | 686,403[117,926] |
| Female | 55 to 59 | 13,280[4,874.8] | 32,769[9,292.6] | 682,902[117,657] |
| Female | 60 to 64 | 11,686[4,295.1] | 28,837[8,191.2] | 600,957[104,082] |
| Female | 65 to 69 | 9,979[3,674.4] | 24,623[7,013.2] | 513,138[89,579] |
| Female | 70 to 74 | 6,975[2,574.9] | NA | 358,639[63,290] |
| Female | 75 to 79 | 4,975[1,843.1] | NA | 255,827[45,777] |
| Female | 80 to 84 | 3,567[1,327.2] | NA | 183,413[33,384] |
| Female | 85 to 89 | 2,470[923.8] | NA | 126,987[23,585] |
| Female | 90 to 94 | 1,271[478.3] | NA | 65,326[12,418] |
| Female | 95+ | 442[167.6] | NA | 22,714[4,433] |
| Total Female |  | 164,746 | 357,961 | 8, 472,359 |
| Male | 15 to 19 | 13,045[4,859.3] | 32,204[9,388.9] | 671,004[122,851] |
| Male | 20 to 24 | 13,577[5,050.2] | 33,518[9,749.3] | 698,389[127,201] |
| Male | 25 to 29 | 13,438[4,992.5] | 33,174[9,630.7] | 691,226[125,367] |
| Male | 30 to 34 | 12,385[4,597.0] | 30,571[8,862.8] | 637,005[115,195] |
| Male | 35 to 39 | 11,443[4,245.8] | 28,247[8,182.8] | 588,586[106,298] |
| Male | 40 to 44 | 10,525[3,905.3] | 25,979[7,525.8] | 541,332[97,821] |
| Male | 45 to 49 | 10,899[4,046.8] | 26,901[7,799.9] | 560,554[101,582] |
| Male | 50 to 54 | 10,990[4,086.3] | 27,127[7,880.3] | 565,253[102,984] |
| Male | 55 to 59 | 10,665[3,973.6] | 26,322[7,670.1] | 548,490[100,747] |
| Male | 60 to 64 | 9,060[3,385.6] | 22,359[6,543.9] | 465,931[86,541] |
| Male | 65 to 69 | 7,516[2,819.4] | 18,548[5,459.5] | 386,508[72,820] |
| Male | 70 to 74 | 5,026[1,894.8] | NA | 258,481[49,556] |
| Male | 75 to 79 | 3,339[1,266.1] | NA | 171,691[33,602] |
| Male | 80 to 84 | 2,139[816.7] | NA | 109,971[22,038] |
| Male | 85 to 89 | 1,240[477.3] | NA | 63,753[13,120] |
| Male | 90 to 94 | 498[193.6] | NA | 25,617[5,428] |
| Male | 95 + | 125[49.0] | NA | 6,416[1,403] |
| Total Male |  | 135,910 | 304,951 | 6,990,206 |
| Total (Male and Female) |  | 300.65[110.51] | 662.91[188.89] | 15,462,564[2,684,712] |

#### Direct costs

Trends of undiscounted excess direct costs due to suboptimal asthma control are shown in ***Figure (3-A)***. In 2019, these costs will be $14.62 billion; this value will increase to $15.08 billion in 2028 and to $15.23 billion in 2038, representing a 4.1% growth during 20 years. Over this period, total direct costs associated with suboptimal asthma control will be $300.65 billion (95%CI 190.1 billion – $411.1 billion).

**Figure 3.**
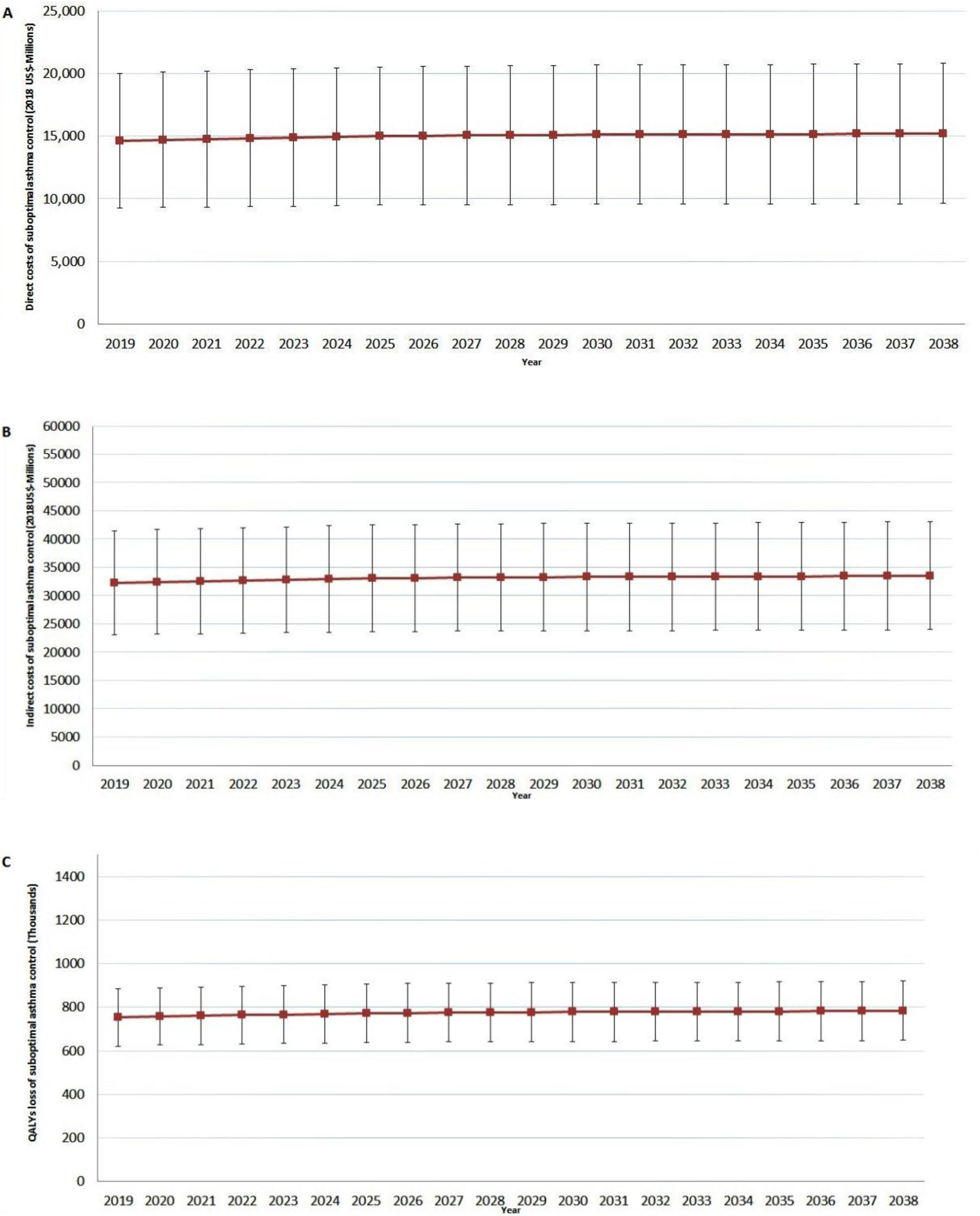
Future trends of undiscounted direct costs (A), undiscounted indirect costs (B) and QALYs loss (C) due to sub-optimal control of asthma in the United State. Squares are point estimates and lines are confidence intervals.

#### Indirect costs

*Figure (3-B)* depicts the projections for indirect costs associated with uncontrolled asthma. These costs will total $32.2 billion in 2019; this value will increase to $33.2 billion in 2028 and to $33.5 billion in 2038, corresponding to a 4.1% increase over 20 years. Over this period, total excess indirect costs associated with suboptimal asthma control will be $662.9 billion (95%CI $474billion – $851billion) over the 20 years.

#### QALYs lost

The total undiscounted QALYs lost due to uncontrolled asthma will be 752,230 in 2019, increasing to 775,791 in 2028 and to 783,474 in 2038; this represents a 4.1% increase during the next 20 years. Over the 20-year study period asthma patients will lose 15.46 million (95%CI 12.77 – 18.14) QALYs due to suboptimal asthma control. Trends of QALYs lost associated with suboptimal asthma control is shown in ***Figure (3-C)***.

### Sensitivity analyses

When future values were discounted at the rate of 3 percent, projections of the burden of suboptimal asthma control changed as follows: total costs decreased to $210.4 billion (95%CI $133 billion – $287.7 billion), total indirect costs decreased to $463.9 billion (95%CI $331.7 billion – $596.1 billion), and total QALYs lost decreased to 10.82 million (95%CI 8.94 – 12.70 million).

### State-level analysis

Results of state-level analyses are provided in *Figure 4* and in Supplementary Materials (Table E8). We divided the total burden over the projected population size for each state to estimate the average projected ‘per capita’ burden of asthma over 20 years. On this metric, Hawaii ranked the first in terms of the direct per-capita costs of suboptimal asthma control ($1,401), while Connecticut ranked the first in terms of indirect per-capita costs ($4,771). Arkansas had the lowest direct and indirect per-capita costs ($666 and $1,543, respectively). In terms of combined direct and indirect costs, Arkansas and Connecticut had the lowest and highest values, respectively ($2,209 and $6,132). Iowa had the lowest per-capita QALYs lost (0.036 QALYs), while New York had the highest values (0.061). The largest increase in direct costs of suboptimal asthma control is expected in South Dakota in which costs are predicted to increase by 14.7% during the projection period, while the smallest increase will be in Georgia (10.2%). Connecticut and D.C. show the largest and smallest increase in per-capita indirect costs of suboptimal asthma control, at 5.8% and 4.7%, respectively.

**Figure 4.**
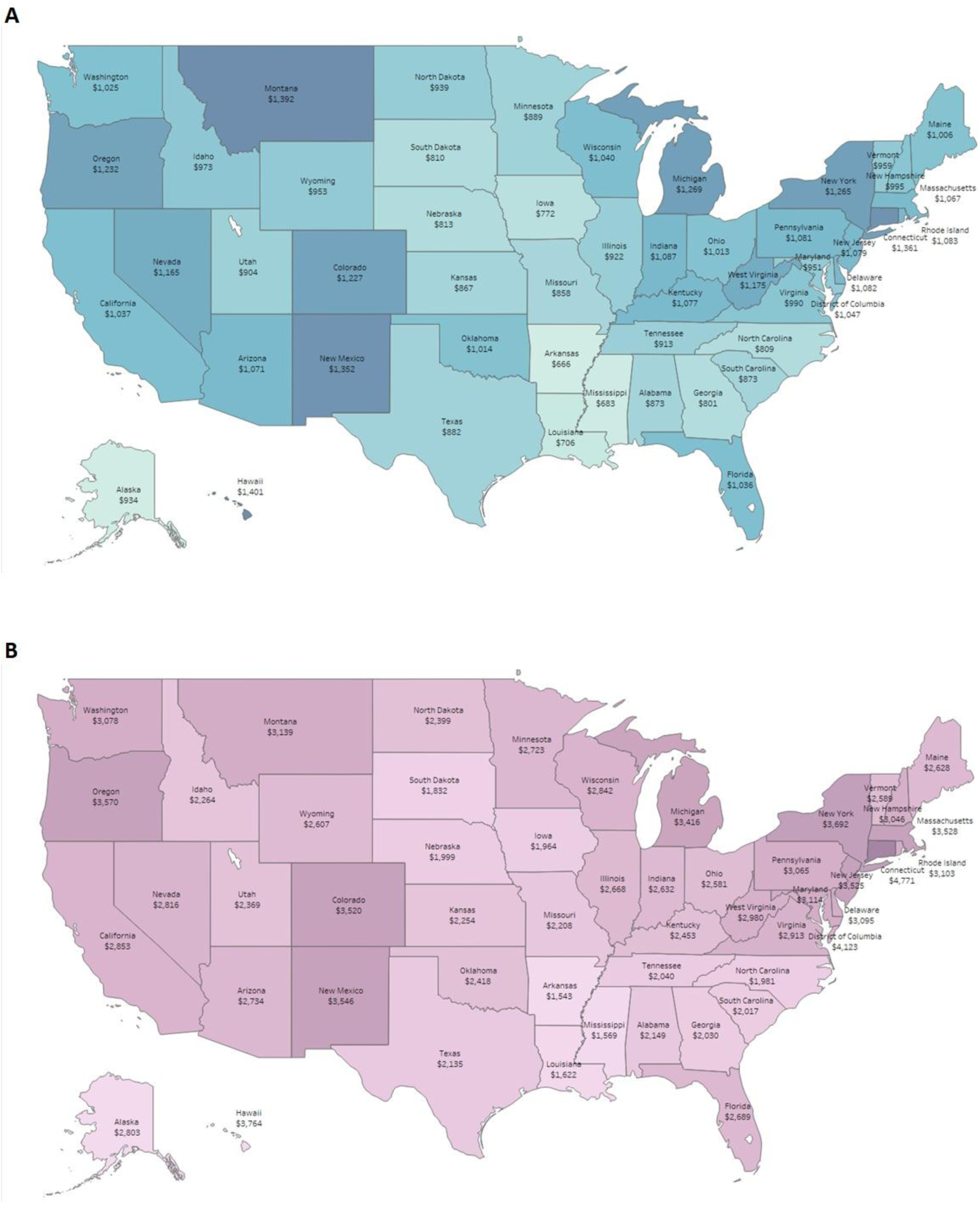
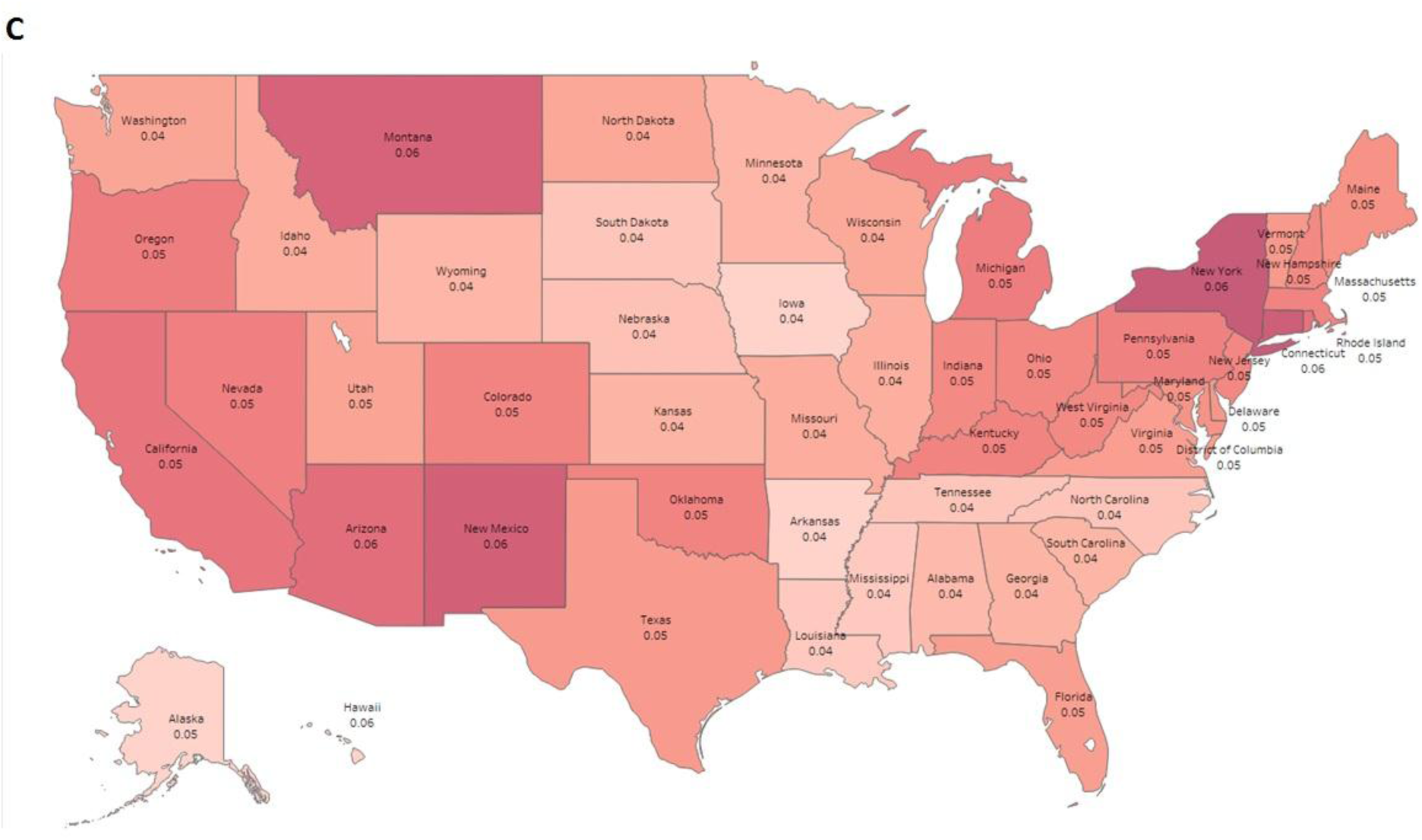
Average 20-year, per-capita estimates of direct costs (A), indirect costs (B), and QALYs lost (C) associated with suboptimal asthma control for each state (2018 US$).All costs were divided to the state’s population to estimate per capita costs.

## Discussion

We predicted the overall burden of uncontrolled asthma over the 2019 – 2038 period in the U.S. adolescent and adult population, in total and across states, if no paradigm shift occurs in the contemporary asthma management. Of the 175.3 million patient-years with asthma in the next 20 years, 52% will be associated with suboptimal asthma control. Total undiscounted direct costs of asthma, across all levels of control, will be $1,537 billion during this period; however, if all patients achieve asthma control during the next 20 years, $300.6 billion in direct costs can be saved. Our results therefore indicate that around 20% of direct costs of asthma can potentially be prevented by achieving asthma control in the population. When indirect costs are added, the potentially preventable burden of sub-optimally controlled asthma will be $963.5 billion. In addition, there will be $15.46 million QALYs lost due to suboptimal asthma control over this period.

Indeed, strategies and interventions towards better asthma control are likely to be associated with costs, and are unlikely to result in complete asthma control in all patients. As such, these values can be seen as population-based estimates of the maximum potential return on investment from strategies that are aimed at improving asthma control. Previous research from the U.S. has repeatedly showed that the prevalence of uncontrolled asthma is remarkably higher than the proportion of patients that fail to achieve asthma control in clinical trials(13–15). Multiple factors are considered as potential culprits for such discrepancy. Such factors include failure to avoid asthma triggers, unacceptably low adherence to controlled medication in asthma patients(9), inefficient uptake of inhaled medications due to poor inhalation techniques(28), over-reliance on reliever versus controller use by both care providers and patients(29), to name a few.

To the best of our knowledge, ours is the first study that provides projections of asthma burden due to suboptimal asthma control. A 2005 study calculated burden of uncontrolled asthma in a managed care setting in U.S., and reported an incremental two-year mean total costs of $7,760 (in 2015 $, equal to incremental one-year costs of $4,873 in 2018 $) between uncontrolled and controlled asthma(30). This estimation is higher than our baseline estimate ($4,699), but nonetheless within the 95%CI of our results. We have previously conducted a similar analysis in the Canadian context using a similar methodology(31). The undiscounted direct and indirect costs (in 2014 Canadian dollars [CAD$]) and QALYs lost attributable to suboptimal asthma control from 2014 to 2033 were, respectively, CAD$24.40 billion, CAD$256.09 billion, and 1.82 million(31). Adjusting for difference in the population sizes (and currency exchange rate for cost values), the corresponding per-capita estimates of these values are $523 (2018 $), $5489.9 (2018 $), and 0.048. The loss of QALY was very similar between the U.S. and the Canadian study (0.04% different). On the other hand, higher per-capita estimate of direct costs for the U.S. is likely due to the differences in healthcare resource utilization and the unit costs of medical services. As for the indirect costs, the overall extent of work impairment is comparable between the two countries (5.07 hours per week in the U.S. vs. 4.10 hours per week in Canada). However, the average weekly income differs in each country, resulting in different estimates of costs due to loss of productivity.

Nurmagambetov et al(32) projected the economic burden of asthma in U.S. at the state level from 2015 to 2020. Total five-year costs associated with asthma ranged from $336.7 million in D.C. to $26.3 billion in California (2014 $)(32). The corresponding per-capita values in (2018 $) ranged from $521 in D.C. to $1,106 in Connecticut. In contrast to Nurmagambetov’s study which reported overall burden of asthma, we reported the excess burden due to suboptimal asthma control. Nevertheless, state-level per-capita estimates of total asthma costs are also obtainable from our model, and are very close to the estimates by Nurmagambetov et al. In the afore-mentioned study, Connecticut had the highest per-capita economic burden of asthma over 5 years(32); our study also highlights this state as having the highest per-capita costs of suboptimal asthma control over 20 years. Similarly, D.C. had the slowest increase rate in the indirect cost of asthma in Nurmagambetov’s study; as well, in our study, it showed the lowest increase in indirect costs due to suboptimal asthma control. Outside North America, our findings about the distribution of asthma control are similar to the trends observed in Europe. For example, we reported that asthma was uncontrolled in 52.13% of patients in the United States, and a similar finding (56.5%) was recently reported by a European study(33).

The strengths of our study include the use of diverse sources of evidence (projection of population growth and aging, prevalence of asthma, distribution of asthma control levels, estimates of resource use and direct and indirect costs and loss of quality of life across control levels). The choice of the analytical framework allowed us to translate such evidence and associated uncertainties into estimates of burden. We conducted multiple systematic reviews and meta-analyses to ensure our estimates of resource use differences across control levels reflect the available evidence. The use of model calibration techniques allowed us to estimate parameters that were not obtainable directly but were estimated such that they remained compatible with evidence studies. For example, we solved for cost values associated with control levels that reflected the differences in healthcare resource use of various types between uncontrolled and controlled asthma, in the meantime adding up to the reported overall costs of asthma from a recent large and representative study(3). The limitations of this study should also be mentioned. Our study assumes the overall prevalence of asthma across age and sex groups will stay the same during the projection period. While such a ‘default’ assumption makes sense for estimating baseline projections for future burden of a disease, it is likely that the contribution of many risk factors (e.g., environmental and occupational pollutions) will change over this time, and the reported ranges (i.e., 95% CIs) in our projections do not reflect this source of uncertainty. Similarly to risk factors, novel therapies will arrive and guidelines and best practice recommendations will change, adding further uncertainty to predictions that are not captured in our results. In addition, the quality of the projections cannot be higher than the quality of the underlying evidence. For example, the observed difference between the burden of uncontrolled and controlled asthma is likely to be confounded by many factors namely the severity of underlying disease. As such, the estimate of the reduction in burden once asthma control is achieved relies on the extent original studies successfully controlled confounding factors. Further, different studies have used different definition of asthma control (e.g., based on cut-offs on the ACT test or symptom control as defined by Global Initiative for Asthma). While we attempted to use a consistent definition of control, the availability of evidence forced us to relax this assumption at times; for example, in estimating the association between outcomes and controlled levels, we considered other definitions of asthma control, such as Global Initiative for Asthma(GINA) and National Asthma Education and Prevention Program (NAEPP) as equivalent of ACT in three studies (8,34,35).

Our findings highlight the sizeable potential for cost saving and improvement in quality of life associated with better asthma control. The conversations about the burden of asthma should focus on the aspects of the burden that can realistically be prevented, rather than just focusing on the overall burden of asthma. As well, research into improving adherence to existing medications should be put on an equal footing with investments in novel asthma therapies. Many of the effective asthma therapies are now off-patent, and research and development in the private sector are understandably shifted towards developing novel therapies. However, healthcare management organizations, patient groups, governments, and society at large, will benefit from investing in areas with proven capacities for improving patient outcomes and reducing costs.

## Acknowledgment

We thank Dr. Zafar Zafari, Zahra Sadat-Fatemi, and Bryan Ng for their help in data collection and comments on the manuscript.

**Authors’ contribution:** MS and JMF conceived the study question. MY developed the analytic plan, performed the literature review, and conducted all the simulations. MS supervised the study progress and provided regular feedback. AA and AS contributed to the study design and performed some of the statistical analyses. MY, wrote the first draft of the manuscript. All authors revised the manuscript and approved the final copy.

## References

1. Nunes C, Pereira AM, Morais-Almeida M. Asthma costs and social impact. Asthma Res Pract. 2017;3:1.

2. CDC - Asthma - Data and Surveillance - Asthma Surveillance Data [Internet]. 2018 [cited 2018 Oct 3]. Available from: https://www.cdc.gov/asthma/asthmadata.htm

3. Nurmagambetov T, Kuwahara R, Garbe P. The Economic Burden of Asthma in the United States, 2008-2013. Ann Am Thorac Soc. 2018 Mar;15(3):348–56.

4. Aalbers R, Vogelmeier C, Kuna P. Achieving asthma control with ICS/LABA: A review of strategies for asthma management and prevention. Respir Med. 2016 Feb 1;111:1–7.

5. 2018 GINA Report, Global Strategy for Asthma Management and Prevention,2018.Available from:www.ginasthma.org.

6. Bateman ED, Bousquet J, Braunstein GL. Is overall asthma control being achieved? A hypothesis-generating study. Eur Respir J. 2001 Apr;17(4):589–95.

7. Doz M, Chouaid C, Com-Ruelle L, Calvo E, Brosa M, Robert J, et al. The association between asthma control, health care costs, and quality of life in France and Spain. BMC Pulm Med. 2013 Mar 22;13:15.

8. Nguyen HV, Nadkarni NV, Sankari U, Mital S, Lye WK, Tan NC. Association between asthma control and asthma cost: Results from a longitudinal study in a primary care setting. Respirol Carlton Vic. 2017;22(3):454–9.

9. Braido F. Failure in Asthma Control: Reasons and Consequences. Scientifica [Internet]. 2013 [cited 2018 Aug 24];2013. Available from: https://www.ncbi.nlm.nih.gov/pmc/articles/PMC3881662/

10. Lozano P, Sullivan SD, Smith DH, Weiss KB. The economic burden of asthma in US children: Estimates from the National Medical Expenditure Survey. J Allergy Clin Immunol. 1999 Nov 1;104(5):957–63.

11. Society ER. “International ERS/ATS guidelines on definition, evaluation and treatment of severe asthma.” Kian Fan Chung, Sally E. Wenzel, Jan L. Brozek, Andrew Bush, Mario Castro, Peter J. Sterk, Ian M. Adcock, Eric D. Bateman, Elisabeth H. Bel, Eugene R. Bleecker, Louis-Philippe Boulet, Christopher Brightling, Pascal Chanez, Sven-Erik Dahlen, Ratko Djukanovic, Urs Frey, Mina Gaga, Peter Gibson, Qutayba Hamid, Nizar N. Jajour, Thais Mauad, Ronald L. Sorkness and W. Gerald Teague. Eur Respir J 2014; 43: 343–373. Eur Respir J. 2018 Jul 1;52(1):1352020.

12. Nathan RA, Sorkness CA, Kosinski M, Schatz M, Li JT, Marcus P, et al. Development of the asthma control test: a survey for assessing asthma control. J Allergy Clin Immunol. 2004 Jan;113(1):59–65.

13. Lee LK, Obi E, Paknis B, Kavati A, Chipps B. Asthma control and disease burden in patients with asthma and allergic comorbidities. J Asthma Off J Assoc Care Asthma. 2018 Feb;55(2):208–19.

14. Williams SA, Wagner S, Kannan H, Bolge SC. The association between asthma control and health care utilization, work productivity loss and health-related quality of life. J Occup Environ Med. 2009 Jul;51(7):780–5.

15. Vietri J, Burslem K, Su J. Poor Asthma control among US workers: health-related quality of life, work impairment, and health care use. J Occup Environ Med. 2014 Apr;56(4):425–30.

16. Stanford RH, Gilsenan AW, Ziemiecki R, Zhou X, Lincourt WR, Ortega H. Predictors of uncontrolled asthma in adult and pediatric patients: analysis of the Asthma Control Characteristics and Prevalence Survey Studies (ACCESS). J Asthma Off J Assoc Care Asthma. 2010 Apr;47(3):257–62.

17. Guilbert TW, Garris C, Jhingran P, Bonafede M, Tomaszewski KJ, Bonus T, et al. Asthma that is not well-controlled is associated with increased healthcare utilization and decreased quality of life. J Asthma Off J Assoc Care Asthma. 2011 Mar;48(2):126–32.

18. Zahran HS, Bailey CM, Qin X, Moorman JE. Assessing asthma control and associated risk factors among persons with current asthma - findings from the child and adult Asthma Call-back Survey. J Asthma Off J Assoc Care Asthma. 2015 Apr;52(3):318–26.

19. Mosnaim G, Lee LK, Carpinella C, Ariely R, Gabriel S, Lugogo NL. The impact of uncontrolled asthma on quality of life among treated, adherent patients with persistent asthma. J Allergy Clin Immunol. 2018 Feb 1;141(2):AB222.

20. Population Projections [Internet]. [cited 2018 Oct 27]. Available from: https://www.census.gov/programs-surveys/popproj.html

21. Colby SL, Ortman JM. Projections of the size and composition of the US population: 2014 to 2060: Population estimates and projections. 2017;

22. Vos T, Abajobir AA, Abate KH, Abbafati C, Abbas KM, Abd-Allah F, et al. Global, regional, and national incidence, prevalence, and years lived with disability for 328 diseases and injuries for 195 countries, 1990–2016: a systematic analysis for the Global Burden of Disease Study 2016. The Lancet. 2017 Sep 16;390(10100):1211–59.

23. Institute for Health Metrics and Evaluation (IHME). GBD Compare. Seattle, WA: IHME, University of Washington, 2015. Available from http://vizhub.healthdata.org/gbd-compare.

24. Highlights of Women’s Earnings in 2012. 2013;91.

25. Mokdad AH, Ballestros K, Echko M, Glenn S, Olsen HE, Mullany E, et al. The State of US Health, 1990-2016: Burden of Diseases, Injuries, and Risk Factors Among US States. JAMA. 2018 Apr 10;319(14):1444–72.

26. Sanders GD, Neumann PJ, Basu A, Brock DW, Feeny D, Krahn M, et al. Recommendations for Conduct, Methodological Practices, and Reporting of Cost-effectiveness Analyses: Second Panel on Cost-Effectiveness in Health and Medicine. JAMA. 2016 Sep 13;316(10):1093–103.

27. US Inflation Calculator [Internet]. US Inflation Calculator. [cited 2018 Dec 11]. Available from: https://www.usinflationcalculator.com/

28. Lavorini F. The Challenge of Delivering Therapeutic Aerosols to Asthma Patients [Internet]. International Scholarly Research Notices. 2013 [cited 2018 Dec 11]. Available from: https://www.hindawi.com/journals/isrn/2013/102418/

29. Shahidi N, FitzGerald JM. Current recommendations for the treatment of mild asthma. J Asthma Allergy. 2010 Dec 8;3:169–76.

30. Sullivan SD. The burden of uncontrolled asthma on the U.S. health care system. Manag Care Langhorne Pa. 2005 Aug;14(8 Suppl):4–7; discussion 25-27.

31. Zafari Z, Sadatsafavi M, Chen W, FitzGerald JM. The projected economic and health burden of sub-optimal asthma control in Canada. Respir Med. 2018;138:7–12.

32. Nurmagambetov T, Khavjou O, Murphy L, Orenstein D. State-level medical and absenteeism cost of asthma in the United States. J Asthma Off J Assoc Care Asthma. 2017 May;54(4):357–70.

33. Braido F, Brusselle G, Guastalla D, Ingrassia E, Nicolini G, Price D, et al. Determinants and impact of suboptimal asthma control in Europe: The INTERNATIONAL CROSS-SECTIONAL AND LONGITUDINAL ASSESSMENT ON ASTHMA CONTROL (LIAISON) study. Respir Res. 2016 May 14;17(1):51.

34. Gold LS, Smith N, Allen-Ramey FC, Nathan RA, Sullivan SD. Associations of patient outcomes with level of asthma control. Ann Allergy Asthma Immunol Off Publ Am Coll Allergy Asthma Immunol. 2012 Oct;109(4):260-265.e2.

35. Sadatsafavi M, McTaggart-Cowan H, Chen W, Mark FitzGerald J, Economic Burden of Asthma (EBA) Study Group. Quality of Life and Asthma Symptom Control: Room for Improvement in Care and Measurement. Value Health J Int Soc Pharmacoeconomics Outcomes Res. 2015 Dec;18(8):1043–9.

