## Supplementary Material for "The projected economic and health burden of uncontrolled asthma in the United States"

#### 1) Input parameters

##### 1.1) Population growth and aging

Estimates of population growth and aging were based on data from two sets of National Population Projections based on the 2010 Census, which were released in 2014 and 2017 by the Census Bureau, Population Division (1)(2). These series used the cohort-component method and historical trends in births, deaths, and international migration, to project the future size and sex and age structure of the national population. We considered the midpoint of these two set of national projection as base case estimate and calculated the standard error around the midpoint (Table E1). Data on state-level population stratified by age and sex were derived from Suburban Stats (3) that provided population information and statistics from each state in the US

***Table E1***. Projected 5-Year Age Groups and Sex Composition of the U.S adolescent and adult (≥14) Population (2016-2040)

| Age | Sex | Population | 2016-2020 | SE | 2020-2025 | SE | 2025-2030 | SE | 2030-2035 | SE | 2035-2040 | SE |
|---|---|---|---|---|---|---|---|---|---|---|---|---|
| **15-19 years** | **Male** | 10,802,000 | -0.0014 | 0.0004 | 0.0018 | 0.0002 | -0.0018 | 0.0006 | 0.0050 | 0.0013 | 0.0043 | 0.0002 |
| **20-24 years** | **Male** | 11,491,000 | -0.0074 | 0.0013 | -0.0003 | 0.0002 | 0.0022 | 0.0002 | -0.0013 | 0.0006 | 0.0051 | 0.0013 |
| **25-29 years** | **Male** | 11,631,000 | 0.0115 | 0.0045 | -0.0064 | 0.0008 | 0.0003 | 0.0002 | 0.0025 | 0.0002 | -0.0008 | 0.0006 |
| **30-34 years** | **Male** | 10,968,000 | 0.0166 | 0.0028 | 0.0114 | 0.0004 | -0.0057 | 0.0007 | 0.0006 | 0.0002 | 0.0028 | 0.0002 |
| **35-39 years** | **Male** | 10,376,000 | 0.0173 | 0.0047 | 0.0143 | 0.0006 | 0.0115 | 0.0003 | -0.0053 | 0.0007 | 0.0009 | 0.0002 |
| **40-44 years** | **Male** | 9,776,000 | 0.0078 | 0.0026 | 0.0160 | 0.0006 | 0.0144 | 0.0005 | 0.0116 | 0.0003 | -0.0049 | 0.0007 |
| **45-49 years** | **Male** | 10,376,000 | -0.0093 | 0.0021 | 0.0043 | 0.0009 | 0.0162 | 0.0006 | 0.0146 | 0.0005 | 0.0118 | 0.0003 |
| **50-54 years** | **Male** | 10,730,000 | -0.0172 | 0.0026 | -0.0064 | 0.0007 | 0.0048 | 0.0009 | 0.0165 | 0.0006 | 0.0148 | 0.0005 |
| **55-59 years** | **Male** | 10,683,000 | -0.0004 | 0.0022 | -0.0151 | 0.0008 | -0.0058 | 0.0006 | 0.0053 | 0.0009 | 0.0169 | 0.0006 |
| **60-64 years** | **Male** | 9,316,000 | 0.0221 | 0.0041 | 0.0015 | 0.0001 | -0.0141 | 0.0007 | -0.0050 | 0.0006 | 0.0060 | 0.0009 |
| **65-69 years** | **Male** | 7,937,000 | 0.0221 | 0.0082 | 0.0207 | 0.0000 | 0.0025 | 0.0000 | -0.0130 | 0.0006 | -0.0042 | 0.0006 |

| | | | | | | | | | | | | |
|---|---|---|---|---|---|---|---|---|---|---|---|---|
| **70-74 years** | **Male** | 5,454,000 | 0.0542 | 0.0051 | 0.0228 | 0.0005 | 0.0216 | 0.0001 | 0.0035 | 0.0000 | -0.0117 | 0.0007 |
| **75-79 years** | **Male** | 3,724,000 | 0.0470 | 0.0054 | 0.0467 | 0.0006 | 0.0240 | 0.0004 | 0.0227 | 0.0001 | 0.0047 | 0.0000 |
| **80-84** | **Male** | 2,453,000 | 0.0305 | 0.0033 | 0.0419 | 0.0005 | 0.0482 | 0.0005 | 0.0256 | 0.0003 | 0.0243 | 0.0001 |
| **85-89** | **Male** | 1,463,000 | 0.0132 | 0.0035 | 0.0289 | 0.0003 | 0.0443 | 0.0004 | 0.0502 | 0.0005 | 0.0279 | 0.0004 |
| **90-94** | **Male** | 605,000 | 0.0317 | 0.0041 | 0.0165 | 0.0005 | 0.0329 | 0.0001 | 0.0475 | 0.0004 | 0.0534 | 0.0007 |
| **95+ years** | **Male** | 156,000 | 0.0665 | 0.0095 | 0.0329 | 0.0008 | 0.0216 | 0.0008 | 0.0377 | 0.0004 | 0.0520 | 0.0005 |
| **15-19 years** | **Female** | 10,328,000 | -0.0002 | 0.0004 | 0.0016 | 0.0001 | -0.0022 | 0.0008 | 0.0052 | 0.0010 | 0.0043 | 0.0001 |
| **20-24 years** | **Female** | 10,890,000 | -0.0057 | 0.0019 | 0.0006 | 0.0001 | 0.0020 | 0.0000 | -0.0018 | 0.0007 | 0.0053 | 0.0009 |
| **25-29 years** | **Female** | 11,259,000 | 0.0082 | 0.0047 | -0.0053 | 0.0006 | 0.0010 | 0.0001 | 0.0023 | 0.0000 | -0.0013 | 0.0007 |
| **30-34 years** | **Female** | 10,818,000 | 0.0126 | 0.0019 | 0.0087 | 0.0008 | -0.0048 | 0.0005 | 0.0013 | 0.0001 | 0.0025 | 0.0000 |
| **35-39 years** | **Female** | 10,397,000 | 0.0152 | 0.0042 | 0.0107 | 0.0006 | 0.0088 | 0.0008 | -0.0044 | 0.0005 | 0.0015 | 0.0001 |
| **40-44 years** | **Female** | 9,920,000 | 0.0059 | 0.0034 | 0.0142 | 0.0003 | 0.0108 | 0.0005 | 0.0088 | 0.0007 | -0.0041 | 0.0004 |
| **45-49 years** | **Female** | 10,572,000 | -0.0088 | 0.0021 | 0.0027 | 0.0004 | 0.0143 | 0.0003 | 0.0109 | 0.0005 | 0.0089 | 0.0007 |
| **50-54 years** | **Female** | 11,109,000 | -0.0193 | 0.0032 | -0.0060 | 0.0006 | 0.0030 | 0.0004 | 0.0145 | 0.0002 | 0.0111 | 0.0004 |
| **55-59 years** | **Female** | 11,297,000 | -0.0012 | 0.0022 | -0.0171 | 0.0007 | -0.0055 | 0.0006 | 0.0033 | 0.0004 | 0.0147 | 0.0001 |
| **60-64 years** | **Female** | 10,167,000 | 0.0203 | 0.0044 | 0.0006 | 0.0003 | -0.0164 | 0.0006 | -0.0051 | 0.0006 | 0.0038 | 0.0004 |
| **65-69 years** | **Female** | 8,883,000 | 0.0234 | 0.0089 | 0.0188 | 0.0005 | 0.0012 | 0.0003 | -0.0156 | 0.0007 | -0.0045 | 0.0007 |
| **70-74 years** | **Female** | 6,356,000 | 0.0527 | 0.0047 | 0.0238 | 0.0008 | 0.0194 | 0.0005 | 0.0019 | 0.0003 | -0.0147 | 0.0007 |
| **75-79 years** | **Female** | 4,644,000 | 0.0442 | 0.0048 | 0.0451 | 0.0006 | 0.0246 | 0.0007 | 0.0203 | 0.0004 | 0.0028 | 0.0002 |
| **80-84** | **Female** | 3,412,000 | 0.0229 | 0.0011 | 0.0391 | 0.0004 | 0.0461 | 0.0006 | 0.0257 | 0.0007 | 0.0214 | 0.0003 |
| **85-89** | **Female** | 2,422,000 | -0.0008 | 0.0001 | 0.0218 | 0.0002 | 0.0411 | 0.0004 | 0.0479 | 0.0005 | 0.0275 | 0.0007 |
| **90-94** | **Female** | 1,278,000 | 0.0116 | 0.0018 | 0.0041 | 0.0000 | 0.0259 | 0.0002 | 0.0443 | 0.0002 | 0.0508 | 0.0006 |
| **95+ years** | **Female** | 456,000 | 0.0463 | 0.0085 | 0.0169 | 0.0001 | 0.0108 | 0.0000 | 0.0317 | 0.0007 | 0.0491 | 0.0003 |

**Source: Bureau UC. Population Projections, [cited 2018 Aug 7]. Available from: https://www.census.gov/programs-surveys/popproj.html**

### 1.2) Sex- and age-specific prevalence of asthma

Estimates of the prevalence of asthma stratified by age and sex for each state were based on the Global Burden of Disease (GBD) studies (4)(5). GBD used the systematic analysis of published studies and available data sources which providing information on prevalence such as the National Health and Nutrition Examination Surveys(5). In these studies, the prevalence of diseases among individuals grouped by age, sex, year, and states were estimated using a range of updated and standardized analytical procedures. More specifically, a Bayesian meta-regression tool was used to determine prevalence and incidence of diseases including asthma (6). The estimated prevalence of asthma in the U.S. population in 2016 served as the baseline for our analysis. We assumed that the prevalence and incidence of asthma within each age and sex group remained constant over time (Table E2).

***Table E2***. Age and sex specific of asthma prevalence in the United States per 100,000

| Age | Sex | Value | Lower 95%CI bound | Upper 95%CI bound |
|---|---|---|---|---|
| 15-19 years | **Male** | **4327** | **4002.2** | **4678.3** |
| 20-24 years | **Male** | **2896** | **2691.5** | **3136.8** |
| 25-29 years | **Male** | **2702** | **2550.3** | **2893.2** |
| 30-34 years | **Male** | **2881** | **2683.8** | **3106.7** |
| 35-39 years | **Male** | **2977** | **2805.2** | **3145.6** |
| 40-44 years | **Male** | **2992** | **2798.9** | **3197.6** |
| 45-49 years | **Male** | **2936** | **2762.6** | **3101.8** |
| 50-54 years | **Male** | **2905** | **2688.3** | **3109.5** |
| 55-59 years | **Male** | **3158** | **2986.0** | **3337.7** |
| 60-64 years | **Male** | **3540** | **3291.8** | **3786.1** |
| 65-69 years | **Male** | **4033** | **3815.6** | **4276.9** |
| 70-74 years | **Male** | **4411** | **4088.8** | **4758.1** |
| 75-79 years | **Male** | **4333** | **4083.6** | **4577.2** |
| 80-84 | **Male** | **4115** | **3792.8** | **4413.1** |
| 85-89 | **Male** | **3983** | **3704.7** | **4260.6** |
| 90-94 | **Male** | **3887** | **3611.5** | **4194.2** |
| 95+ years | **Male** | **3801** | **3380.0** | **4308.4** |
| 15-19 years | **Female** | **5129** | **4798.6** | **5511.9** |
| 20-24 years | **Female** | **4258** | **3967.1** | **4568.4** |
| 25-29 years | **Female** | **4176** | **3946.4** | **4440.4** |
| 30-34 years | **Female** | **4508** | **4227.4** | **4806.4** |
| 35-39 years | **Female** | **4846** | **4603.4** | **5105.6** |
| 40-44 years | **Female** | **5118** | **4828.3** | **5418.0** |
| 45-49 years | **Female** | **5252** | **4987.5** | **5511.3** |
| 50-54 years | **Female** | **5345** | **4998.2** | **5661.4** |
| 55-59 years | **Female** | **5678** | **5403.0** | **5976.7** |
| 60-64 years | **Female** | **6159** | **5764.7** | **6547.1** |
| 65-69 years | **Female** | **6772** | **6442.3** | **7138.1** |
| 70-74 years | **Female** | **7040** | **6565.3** | **7536.8** |
| 75-79 years | **Female** | **6289** | **5950.4** | **6637.6** |
| 80-84 | **Female** | **5313** | **4907.4** | **5680.4** |
| 85-89 | **Female** | **4799** | **4463.4** | **5105.4** |
| 90-94 | **Female** | **4458** | **4170.1** | **4774.0** |
| 95+ years | **Female** | **4181** | **3697.9** | **4748.6** |

**Source: Institute for Health Metrics and Evaluation (IHME). GBD Compare, Seattle, WA: IHME, University of Washington, 2015. Available from http://vizhub.healthdata.org/gbd-compare**

**1.3) Distributions of asthma control levels and its association with sex and age**

The distribution of levels of asthma control within a given sex and age group was derived using calibration techniques from a recent study based on U.S. National Health and Wellness Survey (NHWS) by Lee et al (7). This study was based on the data from 1,923 patients from NHWS between 2011 and 2013. The NHWS is a representative, large-scale survey of the adult population (≥18 years), and assesses health status and outcomes across a wide array of diseases. In particular, persons with asthma are categorized based on their score on the Asthma Control Test. Possible scores are very poorly controlled (scores ≤ 15; 29.3%), not well controlled (score 16-19; 25.1%), or well controlled (score 20-25; 45.7%). ACT is a validated and widely used instrument to measure asthma control; studies have demonstrated that ACT is reliable, valid, and responsive to changes in asthma control over time (8,9). Lee et.al did not provide direct estimates of the prevalence of asthma control within sex and age groups. However, this information could be estimated indirectly from the reported proportion of asthma patients falling into each of three control categories, as well as the sex and age distribution of the sample within each control category. This information was sufficient to estimate the coefficient of the following multinomial logit equations:

Probability of not well-controlled = $\frac{\exp(\beta_0+\beta_1.age+\beta_2.sex)}{1+\exp(\beta_0+\beta_1.age+\beta_2.sex)+\exp(\beta_3+\beta_4.age+\beta_5.sex)}$

Probability of well-controlled = $\frac{\exp(\beta_3+\beta_4.age+\beta_5.sex)}{1+\exp(\beta_0+\beta_1.age+\beta_2.sex)+\exp(\beta_3+\beta_4.age+\beta_5.sex)}$

Probability of very poorly controlled = 1- (Probability of not well-controlled + Probability of well-controlled)

After estimating probability of each of the three levels of control, we merged “very poorly control” and “not-well control” into “uncontrolled group “.

Uncertainty around the estimated coefficients as well as their covariance were computed through parametric bootstrapping, by randomly sampling variables representing calibration targets from their reported distributions and repeating the calibration process 1,000 times. Covariance matrix of coefficients are provided in (Table E3)

***Table E3***: Covariance matrix of coefficients in multinomial logit equations

| | **β 1** | **β 2** | **β 3** | **β 4** | **β 5** | **β 6** |
|---|---|---|---|---|---|---|
| **β 1** | 0.07069 | -0.00071 | -0.02307 | 0.02611 | -0.00026 | -0.00992 |
| **β 2** | -0.00071 | 0.00002 | -0.00005 | -0.00030 | 0.00001 | -0.00004 |
| **β 3** | -0.02307 | -0.00005 | 0.02191 | -0.00772 | -0.00008 | 0.01043 |
| **β 4** | 0.02611 | -0.00030 | -0.00772 | 0.04756 | -0.00064 | -0.01250 |
| **β 5** | -0.00026 | 0.00001 | -0.00008 | -0.00064 | 0.00002 | -0.00009 |
| **β 6** | -0.00992 | -0.00004 | 0.01043 | -0.01250 | -0.00009 | 0.01757 |

### 1.4) Costs and health differences across control levels

#### 1.4.1) Literature review and meta-analysis

We conducted a literature review to retrieve all relevant studies that report on the association between the level of asthma control on healthcare resource utilization, overall work impairment, and health-related quality of life, adjusting for potential confounding variables. The data from these studies were used to calculate the total direct costs, indirect costs, and QALYs lost as a result of sub-optimal asthma control in the Unites States, as described I the main text. All relevant studies, up to September 2018, were retrieved from MEDLINE via Ovid using a search strategy designed with the help of a librarian (Table E4). Two reviewers [MY-BN] independently screened all titles and abstracts retrieved during the initial search on PubMed. The full texts were obtained for each study that was deemed potentially relevant. They were screened against the inclusion criteria outlined below. Any disagreement concerning the eligibility of a study for this review was resolved through discussion with the third reviewer (MS). Studies were included in the rapid literature review if they evaluated the adjusted association between asthma control and at least one of the selected outcomes in the US. Outcomes of interest were adjusted means or odds ratios (95% CIs) associated with the use of healthcare utilization across control levels, overall work impairment including both absenteeism and presenteeism by control levels (10), and health-related quality of life across control levels. We excluded studies that were performed outside of the United States, as well as paediatric asthma studies. The study selection process is presented in a Preferred Reporting Items for Systematic Review and Meta-Analysis (PRISMA) flow chart (Figure E1).The meta-analysis was performed using Stata (version 14). The heterogeneity of studies in the meta-analysis was assessed using the Q test to quantify heterogeneity.

***Table E4*: **Search strategy

| PUBMED | | |
|---|---|---|
| **No** | **Search Term** | **Result** |
| 1 | (Asthma* OR Anti-asthmatic or Bronchial Hyperreactivity OR Respiratory Hypersensitivity or Reactive airway*) | 247784 |
| 2 | ((Controlled) OR Uncontrolled) OR Well controlled | 1184416 |
| 3 | 1 AND 2 | 29028 |
| 4 | ((((healthcare utilization) OR hospitalization) OR emergency visit) OR medication) OR direct cost | 1319845 |
| 5 | ((((work impairment) OR productivity loss) OR absenteeism) OR presenteeism') OR indirect cost | 50148 |
| 6 | (HRQol) OR quality of life | 337519 |
| 7 | 4 OR 5 OR 6 | 1657204 |
| 8 | 3 AND 7 | 10841 |
| 9 | 8 AND United States[PL] | 3615 |

***Figure E1: Process of selection of studies for systematic review based on PRISMA flow diagram***

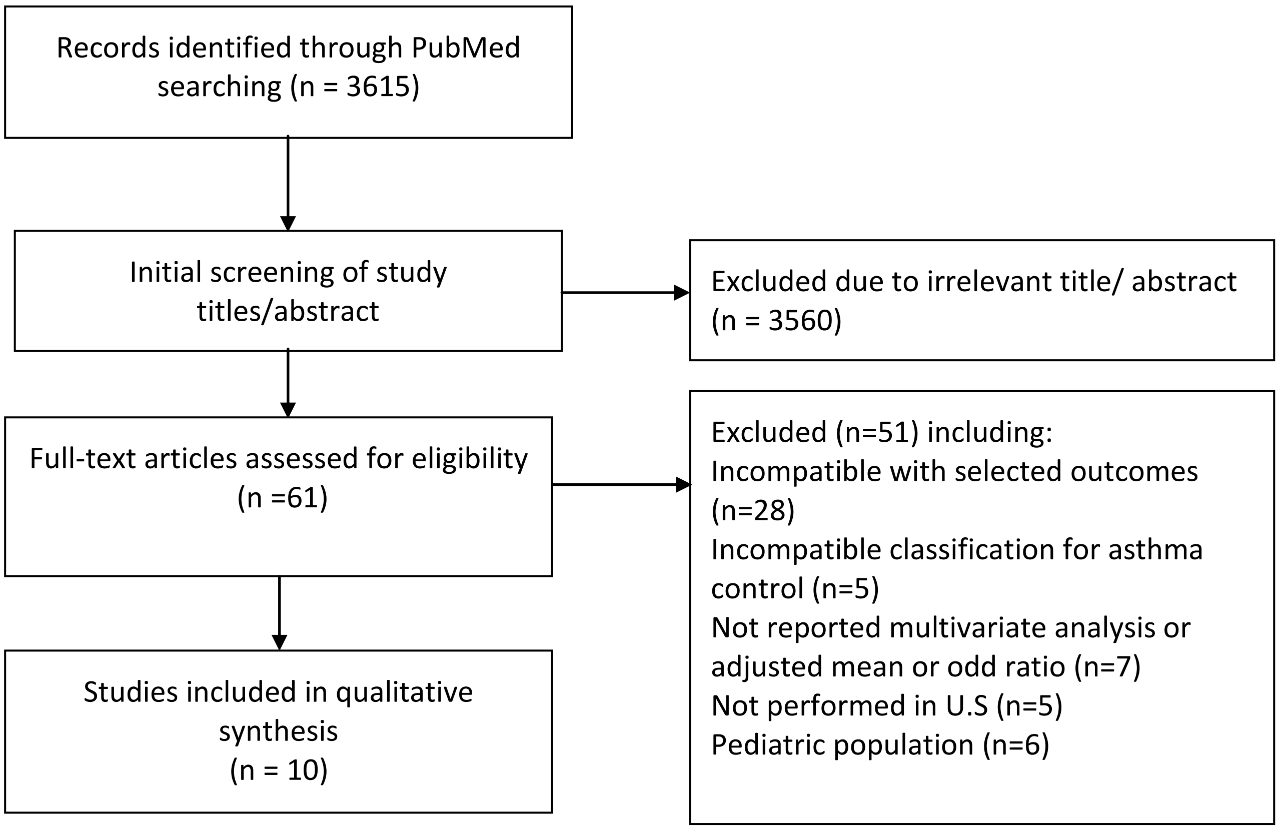

#### 1.4.2) Meta-Analysis

##### 1.4.2.1) Health care provider visits

Five studies reported on this outcome (11–15), with a combined sample size of 13,135 asthma patients, including 5,969 well-controlled and 7,166 uncontrolled asthma were examined to estimate adjusted OR (95% CIs) associated with health care provider visit across level of control. A random-effects model was used to estimate the pooled adjusted OR (95% CIs) associated with health care provider visits in uncontrolled group versus well-controlled. The pooled estimate of the odd ratio across groups was 1.86 (95% CI 1.34 to 2.38; P<0.001).

##### 1.4.2.2) Emergency visits

Five studies (11,12,16–18) with a combined sample size of 14,003 were included. The sample consisted of 7,263 well-controlled and 6,741 uncontrolled asthma; the samples were examined to estimate adjusted OR (95% CIs) associated with emergency visit across level of control. A random-effects model was used to estimate the pooled adjusted OR (95% CIs) associated with emergency visits in uncontrolled group versus well-controlled. The pooled estimate of the odd ratio across groups was 1.44 (95% CI 1.39 to 1.49; P<0.001).

##### 1.4.2.3) Hospitalization

Three studies (11–13) reported on the data of a total of 10,198 patients for this outcome, including 4,771 well-controlled and 5,427 uncontrolled asthma who were to estimate adjusted OR (95% CIs) associated with hospitalization across level of control. A random-effects model was used to estimate the pooled adjusted OR (95% CIs) associated with hospitalization in uncontrolled group versus well-controlled. The pooled estimate of the odd ratio across groups was 1.54 (95% CI 1.25 to 1.80; P<0.001).

##### 1.4.2.4) Medication use

Three studies (12,16,19) with a combined sample size of 15,261 were included. The sample consisted of 8,997 well-controlled and 6,264 uncontrolled asthma, who were examined to estimate adjusted OR (95% CIs) associated with controller medication use across level of control. A random-effects model was used to estimate the pooled adjusted OR (95% CIs) associated with medication use in uncontrolled group versus well-controlled. The pooled estimate of the odd ratio across groups was 1.58 (95% CI 1.26 to 1.90; P<0.001).

##### 1.4.2.5) Overall work impairment

Three studies (7,11,13) with a combined sample size of 9,628 were included (including5,011 well-controlled and 4,617 uncontrolled). We subtracted the baseline percent of impairments from the follow-up percent to find the change over time. Next, we subtracted the change in the control group from the corresponding change in the intervention group to quantify the net mean difference of overall work impairment (%). A random-effects model was used to estimate the pooled standardized mean difference in overall work impairment (%) between the uncontrolled and the well-controlled group. The pooled estimate of the standardized mean difference of overall work impairment (%) across groups was 12.70% (95% CI 9.41 to 15.98; P<0.001). Assuming 52 workweeks in year, this translates to a loss of 6.6 weeks of productivity per year (5.07 hours per week) lost for each patient with uncontrolled asthma.

##### 1.4.2.6) Health-Related Quality of Life

Five studies(7,11,13,20,21), with a combined sample size of 10589 were included (with 5,650 having well-controlled and 4,939 having uncontrolled asthma). Three studies (7,13,20) reported adjusted mean SF-6D score in well-controlled and uncontrolled of asthma. William et al (11) reported SF-8 score across level of control. In order to achieve consistency, we predicted SF-6D index score form from the SF-8 based on P.Wang et al (22). Also one study reported EQ-5D score across asthma symptom control in Canada (21) . We included all studies that assessed health-related quality of life using a generic questionnaire and studies which used specific disease questionnaires were excluded from this review. A random-effects model was used to estimate the pooled mean of excessive QALYs loss between the uncontrolled and the well-controlled group. The pooled estimate of the mean difference of health utility score across groups was 0.07 (95% CI 0.06 to 0.09; P<0.001). (Table E1)

***Table E5***: Summary of studies assessing adjusted odd ratio associate with use of health care resource by level of control asthma

| Author | Year | Survey | Classification | Well-Controlled (n) | Uncontrolled(n) | OR | Se |
|---|---|---|---|---|---|---|---|
| **Health care provider Visits** | | | | | | | |
| A. Williams | 2009 | NHWS | ACT* | 2912 | 2676 | 1.76 | 0.02 |
| H. Stanford | 2010 | ACCESS | ACT | 921 | 1317 | 2.37 | 0.57 |
| W .Guilbert | 2011 | HarrisPollOnline | ACT | 277 | 422 | 3.30 | 1.84 |
| S. Gold | 2012 | AIM | GINA† | 638 | 1855 | 5.60 | 1.79 |
| J.Vietri | 2014 | NHWS | ACT | 1221 | 805 | 1.19 | 0.36 |
| **Emergency Visits** | | | | | | | |
| A. Williams | 2009 | NHWS | ACT | 2912 | 2676 | 1.44 | 0.03 |
| K.Nguyen | 2010 | BRFSS | NAEPP‡ | 2215 | 891 | 3.90 | 2.65 |
| W .Guilbert | 2011 | HarrisPollOnline | ACT | 277 | 422 | 11.3 | 25.5 |
| S. Gold | 2012 | AIM | GINA | 638 | 1855 | 2.10 | 0.50 |
| J.Vietri | 2014 | NHWS | ACT | 1221 | 805 | 1.97 | 0.54 |
| **Hospitalization** | | | | | | | |
| A. Williams | 2009 | NHWS | ACT | 2912 | 2676 | 1.45 | 0.06 |
| S. Gold | 2012 | AIM | GINA | 638 | 1855 | 2.20 | 0.61 |
| J.Vietri | 2014 | NHWS | ACT | 1221 | 805 | 2.15 | 0.78 |
| **Medication use** | | | | | | | |
| K.Nguyen | 2010 | BRFSS | NAEPP | 2215 | 891 | 2.60 | 1.33 |
| S. Gold | 2012 | AIM | GINA | 638 | 1855 | 1.20 | 0.31 |
| H.S Zahran | 2015 | BRFSS | ACT | 6144 | 3518 | 1.70 | 0.10 |

**ACT= Asthma Control Test*
*†GINA: Global Initiative for Asthma*
*‡ NAEPP: National Asthma Education and Prevention Program*

***Table E6:*** Summary of studies assessing adjusted mean difference associate with percentage of overall work impairment by level of control asthma

| Author | Year | Survey | Classification | Well-Controlled (n) | Uncontrolled(n) | Mean Difference (%) | Se |
|---|---|---|---|---|---|---|---|
| A. Williams | 2009 | NHWS | ACT | 2912 | 2676 | 10.81 | 1.30 |
| J.Vietri | 2014 | NHWS | ACT | 1221 | 805 | 15.50 | 3.08 |
| K.lee | 2017 | NHWS | ACT | 878 | 1045 | 18.86 | 7.30 |

***Table E7***: Summary of studies assessing adjusted mean difference associate with quality of life by level of control asthma

| Author | Year | Survey | Classification | Well-Controlled (n) | Uncontrolled (n) | Utility instrument | Mean Difference of Utility score | Se |
|---|---|---|---|---|---|---|---|---|
| A. Williams | 2009 | NHWS | ACT | 2912 | 2676 | SF-6D | 0.11 | 0.03 |
| J.Vietri | 2014 | NHWS | ACT | 1221 | 805 | SF-6D | 0.07 | 0.03 |
| M.Sadatsafavi | 2015 | EBA | GINA | 639 | 322 | EQ-5D | 0.05 | 0.01 |
| K.lee | 2017 | NHWS | ACT | 878 | 1045 | SF-6D | 0.09 | 0.005 |
| G.Mansnaim | 2018 | NHWS | ACT | -- | -- | SF-6D | 0.07 | 0.02 |

### 1.5) Direct medical costs

We calculated the annual direct costs of controlled and uncontrolled asthma per-person based on: 1) the annual per-person excess medical costs of asthma (in 2015 US$) estimated by NurmagambetovT et al (23), 2) the estimated prevalence of controlled and uncontrolled asthma, and 3) pooled odds ratio of healthcare utilization associated with uncontrolled versus controlled asthma. A recent economic burden of asthma study in U.S. (2) used data from the 2008-2013 household components of the Medical Expenditure Panel Survey (MEPS) was deemed to be the most reliable source of asthma costs. These estimates totalled $3,266 (in 2015 U.S dollars), of which $1,830 was attributable to prescription medication, $640 to office visits, $529 to hospitalizations, $176 to hospital-based outpatient visits, and $105 to emergency room visits (23). We considered above estimation as total excess costs of healthcare utilization for asthma and based on the following formula, we calculated the annual per-person direct costs of well-controlled asthma in terms of health care utilization. We multiplied these values by the adjusted pooled odds ratio of healthcare utilization in the uncontrolled group compared to the well-controlled group. This allowed us to estimate the excess direct costs of uncontrolled asthma based on healthcare utilization in 2016:

$$C = P.x + (1 - P)x.RR$$

Where

C =Excess cost of healthcare utilization for asthma,

X = Baseline cost of healthcare utilization for asthma,

RR= Adjusted pooled ratio of healthcare utilization between uncontrolled and controlled groups, and

P= Proportion of asthma patients that are controlled.

We applied a probabilistic version of the above equation to calculate the annual per-person direct costs for the year 2016 as the baseline year. The annual per-person excess direct costs of suboptimal asthma control in 2018 US$ were used for all projections pertaining to the U.S. population for the years 2019 to 2038 (Table 1). For state-level analyses, we multiplied our 2016 per-person excess costs by a state-level adjustment factor derived from Nurmagambetov T et al (24).

**1.6) indirect medical costs**

Using the pooled estimate of the standardized mean difference of overall work impairment (%) from presentism and absenteeism between the uncontrolled and controlled groups, we estimated the total excess indirect costs of suboptimal asthma control. To estimate the monetary value of lost productivity, 2012's age- and sex-specific wages reported by the Bureau of Labour Statistics were used (25). These wages were applied to the excess percentage of overall work impairment resulting from uncontrolled asthma. We solved for the equation probabilistically to calculate the annual per-person indirect cost in 2018 US $. For state-level analysis, the median weekly earnings from full-time wages were used as reported for each state and sex (25). These wages were applied to the excess percentage of work impairment resulting from uncontrolled asthma in order to estimate the excess uncontrolled cost of asthma in each state.

### 2) Modeling approach:

The analytical framework for projections was an open-population probabilistic time-in-state model of asthma control. Time-in-state models estimate the progression of a population across mutually exclusive health states in relation to risk factors, without having to identify the transition rates across disease states, making them ideal for the present study. The model stratified U.S. asthma population (≥14 years old) into age groups (with 5-year bands), sex (male and female), and three levels of asthma control as defined based on the Asthma Control Test (26): well-controlled, not well- controlled, and very poorly controlled, creating a total of 240 states. Because the estimates are generated separately for each year, there is no need for consideration of transition matrix and for modeling entrance and exit in the population (e.g., birth or death); such population dynamics are already incorporated through the use of population size and age/sex structure projections as input parameters for the model. For reporting the results, we considered the not-well controlled and very poorly controlled groups together as the sub-optimally controlled asthma. This model structure related the prevalence of asthma and the distribution of asthma control to age and sex. As such, the implicit assumption is that the impact of all other risk factors (e.g. environmental variables) will remain constant throughout the projection period. All analyses were conducted in using R (version 3.2.2) (12).

#### 2.1) Uncertainty analysis

The uncertainty in projections was quantified through Monte Carlo simulation. Uncertainty in each of the underlying modeling components were characterized by assigning probability distribution to point estimates, and the model was run for 10,000 times for baseline estimate in

2016 as well as projection estimation from 2019-2038. Results were reported in terms of 95% confidence interval [CI] around point estimates of projections.

**Table E8) the state-level undiscounted projected 20-years direct cost, indirect cost and QALYs lost associated with suboptimal control of asthma**

| | | (US$ Million) | | | Average Per State Population | | | Increase from 2019 to 2038 (%) | |
|---|---|---|---|---|---|---|---|---|---|
| | **State** | **Excess Direct Costs** | **Excess Indirect Costs** | **Excess QALYs lost** | **Excess Direct Costs(US$)** | **Excess Indirect Costs (US$)** | **Excess QALYs lost** | **Excess direct costs** | **Excess indirect cots** |
| 1 | **Alabama** | 2,657.14 | 7,266.53 | 125,459 | 873.19 | 2149.26 | 0.0412 | 11.49 | 5.11 |
| 2 | **Alaska** | 608.60 | 2,006.47 | 29,765 | 933.53 | 2802.93 | 0.0457 | 12.86 | 5.32 |
| 3 | **Arizona** | 4,195.87 | 11,895.14 | 215,997 | 1071.33 | 2734.29 | 0.0552 | 11.42 | 5.23 |
| 4 | **Arkansas** | 1,297.15 | 3,276.06 | 70,114 | 666.12 | 1543.12 | 0.0360 | 11.95 | 5.08 |
| 5 | **California** | 21,400.50 | 69,785.88 | 1,110,478 | 1036.59 | 2852.72 | 0.0538 | 11.23 | 5.43 |
| 6 | **Colombia** | 658.84 | 2,718.81 | 30,524 | 1046.62 | 4122.61 | 0.0485 | 13.32 | 4.77 |
| 7 | **Colorado** | 3,731.70 | 12,545.14 | 155,159 | 1227.05 | 3520.50 | 0.0510 | 11.34 | 5.51 |
| 8 | **Connecticut** | 3,219.70 | 12,351.85 | 140,869 | 1361.50 | 4771.10 | 0.0596 | 12.19 | 5.83 |
| 9 | **Delaware** | 874.84 | 2,618.36 | 38,602 | 1082.01 | 3095.22 | 0.0477 | 13.43 | 5.29 |
| 10 | **Florida** | 12,099.86 | 33,615.37 | 538,478 | 1035.98 | 2689.01 | 0.0461 | 12.33 | 5.23 |
| 11 | **Georgia** | 4,534.01 | 13,413.05 | 239,828 | 800.74 | 2029.79 | 0.0424 | 10.25 | 5.52 |
| 12 | **Hawaii** | 1,492.96 | 4,267.22 | 63,035 | 1401.05 | 3763.68 | 0.0592 | 13.24 | 5.22 |
| 13 | **Idaho** | 1,098.45 | 2,805.71 | 49,095 | 973.40 | 2263.53 | 0.0435 | 13.27 | 5.17 |
| 14 | **Illinois** | 7,136.70 | 23,201.39 | 334,228 | 922.09 | 2668.14 | 0.0432 | 10.39 | 5.52 |
| 15 | **Indiana** | 4,237.71 | 11,631.51 | 194,255 | 1087.28 | 2632.17 | 0.0498 | 11.84 | 5.32 |
| 16 | **Iowa** | 1,547.23 | 4,261.43 | 72,006 | 771.77 | 1964.07 | 0.0359 | 12.87 | 5.27 |
| 17 | **Kansas** | 1,632.50 | 4,648.05 | 79,541 | 866.55 | 2254.20 | 0.0422 | 11.76 | 5.24 |
| 18 | **Kentucky** | 2,957.52 | 7,553.22 | 140,280 | 1077.29 | 2453.37 | 0.0511 | 11.75 | 5.31 |
| 19 | **Louisiana** | 1,986.11 | 5,160.68 | 106,357 | 706.07 | 1621.51 | 0.0378 | 11.32 | 5.08 |
| 20 | **Maine** | 1,086.61 | 2,977.14 | 52,451 | 1006.32 | 2628.01 | 0.0486 | 13.02 | 5.45 |
| 21 | **Maryland** | 3,474.82 | 12,858.71 | 176,601 | 950.71 | 3114.06 | 0.0483 | 10.77 | 5.57 |
| 22 | **Massachusetts** | 4,360.65 | 16,071.96 | 207,783 | 1067.34 | 3528.26 | 0.0509 | 11.79 | 5.62 |
| 23 | **Michigan** | 7,586.70 | 22,965.44 | 311,699 | 1269.41 | 3415.92 | 0.0522 | 11.70 | 5.35 |
| 24 | **Minnesota** | 2,927.08 | 10,050.20 | 137,579 | 889.24 | 2722.83 | 0.0418 | 11.35 | 5.48 |
| 25 | **Mississippi** | 1,324.55 | 3,369.88 | 73,510 | 682.78 | 1568.99 | 0.0379 | 11.89 | 5.19 |
| 26 | **Missouri** | 3,176.53 | 9,117.72 | 161,441 | 858.20 | 2208.31 | 0.0436 | 11.86 | 5.32 |
| 27 | **Montana** | 1,186.48 | 2,807.18 | 49,332 | 1391.94 | 3138.54 | 0.0579 | 13.42 | 5.06 |
| 28 | **Nebraska** | 1,048.32 | 2,794.94 | 50,934 | 813.18 | 1998.74 | 0.0395 | 13.47 | 5.10 |
| 29 | **Nevada** | 2,117.78 | 5,774.92 | 94,005 | 1165.22 | 2815.68 | 0.0517 | 11.39 | 5.52 |
| 30 | **New Hampshire** | 1,049.13 | 3,457.69 | 52,701 | 994.67 | 3046.44 | 0.0500 | 12.77 | 5.80 |
| 31 | **New Jersey** | 5,844.53 | 21,324.99 | 272,484 | 1078.93 | 3525.31 | 0.0503 | 11.53 | 5.77 |
| 32 | **New Mexico** | 1,986.79 | 5,634.15 | 86,200 | 1351.62 | 3545.64 | 0.0586 | 11.34 | 5.17 |
| 33 | **New York** | 14,683.27 | 48,259.07 | 703,527 | 1264.90 | 3692.27 | 0.0606 | 11.17 | 5.54 |
| 34 | **North Carolina** | 4,597.57 | 12,860.23 | 221,102 | 808.55 | 1980.68 | 0.0389 | 11.34 | 5.37 |
| 35 | **North Dakota** | 622.27 | 1,629.18 | 29,515 | 939.27 | 2399.12 | 0.0446 | 14.37 | 4.88 |

| | | | | | | | | | |
|---|---|---|---|---|---|---|---|---|---|
| 36 | **Ohio** | 7,024.44 | 19,982.04 | 347,806 | 1013.04 | 2581.27 | 0.0502 | 11.68 | 5.36 |
| 37 | **Oklahoma** | 2,412.30 | 6,398.47 | 121,059 | 1013.55 | 2417.88 | 0.0509 | 12.10 | 5.11 |
| 38 | **Oregon** | 3,087.12 | 9,926.95 | 130,343 | 1232.26 | 3570.31 | 0.0520 | 11.71 | 5.13 |
| 39 | **Pennsylvania** | 8,493.65 | 26,001.48 | 402,501 | 1081.05 | 3065.35 | 0.0512 | 11.08 | 5.42 |
| 40 | **Rhode Island** | 972.94 | 2,923.93 | 48,873 | 1083.48 | 3102.93 | 0.0544 | 13.49 | 5.49 |
| 41 | **South Carolina** | 2,570.77 | 6,631.88 | 123,288 | 873.04 | 2017.08 | 0.0419 | 11.20 | 5.00 |
| 42 | **South Dakota** | 591.23 | 1,379.64 | 28,224 | 809.91 | 1831.62 | 0.0387 | 14.77 | 5.02 |
| 43 | **Tennessee** | 3,537.15 | 9,011.06 | 150,322 | 913.30 | 2039.90 | 0.0388 | 12.02 | 5.21 |
| 44 | **Texas** | 12,156.02 | 35,044.33 | 643,652 | 882.42 | 2134.53 | 0.0467 | 10.34 | 5.40 |
| 45 | **Utah** | 1,516.86 | 4,564.83 | 75,829 | 904.21 | 2369.40 | 0.0452 | 11.22 | 5.32 |
| 46 | **Vermont** | 622.70 | 1,729.84 | 30,657 | 958.67 | 2588.77 | 0.0472 | 14.04 | 5.36 |
| 47 | **Virginia** | 4,791.85 | 16,121.81 | 224,010 | 989.95 | 2913.47 | 0.0463 | 10.92 | 5.42 |
| 48 | **Washington** | 4,241.21 | 14,524.67 | 185,250 | 1024.87 | 3077.68 | 0.0448 | 10.78 | 5.39 |
| 49 | **West Virginia** | 1,641.55 | 4,395.39 | 69,820 | 1174.56 | 2979.77 | 0.0500 | 12.33 | 5.12 |
| 50 | **Wisconsin** | 3,681.97 | 11,177.93 | 157,250 | 1039.92 | 2842.20 | 0.0444 | 11.65 | 5.46 |
| 51 | **Wyoming** | 569.70 | 1,618.38 | 25,010 | 953.10 | 2607.16 | 0.0418 | 14.11 | 5.16 |

### 3) References


1. Bureau UC. Population Projections [Internet]. [cited 2018 Aug 7]. Available from: https://www.census.gov/programs-surveys/popproj.html

2. Colby SL, Ortman JM. Projections of the size and composition of the US population: 2014 to 2060: Population estimates and projections. 2017;

3. Stats S. Population Information and Statistics From Every City, State and County in the US [Internet]. Vietri. [cited 2018 Aug 25]. Available from: https://suburbanstats.org/

4. Institute for Health Metrics and Evaluation (IHME). GBD Compare. Seattle, WA: IHME, University of Washington, 2015. Available from http://vizhub.healthdata.org/gbd-compare.

5. Mokdad AH, Ballestros K, Echko M, Glenn S, Olsen HE, Mullany E, et al. The State of US Health, 1990-2016: Burden of Diseases, Injuries, and Risk Factors Among US States. JAMA. 2018 Apr 10;319(14):1444–72.

6. Vos T, Abajobir AA, Abate KH, Abbafati C, Abbas KM, Abd-Allah F, et al. Global, regional, and national incidence, prevalence, and years lived with disability for 328 diseases and injuries for 195 countries, 1990–2016: a systematic analysis for the Global Burden of Disease Study 2016. The Lancet. 2017 Sep 16;390(10100):1211–59.

7. Lee LK, Obi E, Paknis B, Kavati A, Chipps B. Asthma control and disease burden in patients with asthma and allergic comorbidities. J Asthma Off J Assoc Care Asthma. 2018 Feb;55(2):208–19.

8. Schatz M, Sorkness CA, Li JT, Marcus P, Murray JJ, Nathan RA, et al. Asthma Control Test: reliability, validity, and responsiveness in patients not previously followed by asthma specialists. J Allergy Clin Immunol. 2006 Mar;117(3):549–56.

9. Schatz M, Kosinski M, Yarlas AS, Hanlon J, Watson ME, Jhingran P. The minimally important difference of the Asthma Control Test. J Allergy Clin Immunol. 2009 Oct;124(4):719-723.e1.

10. Reilly MC, Zbrozek AS, Dukes EM. The validity and reproducibility of a work productivity and activity impairment instrument. PharmacoEconomics. 1993 Nov;4(5):353–65.

11. Williams SA, Wagner S, Kannan H, Bolge SC. The association between asthma control and health care utilization, work productivity loss and health-related quality of life. J Occup Environ Med. 2009 Jul;51(7):780–5.

12. Gold LS, Smith N, Allen-Ramey FC, Nathan RA, Sullivan SD. Associations of patient outcomes with level of asthma control. Ann Allergy Asthma Immunol Off Publ Am Coll Allergy Asthma Immunol. 2012 Oct;109(4):260-265.e2.

13. Vietri J, Burslem K, Su J. Poor Asthma control among US workers: health-related quality of life, work impairment, and health care use. J Occup Environ Med. 2014 Apr;56(4):425–30.

14. Guilbert TW, Garris C, Jhingran P, Bonafede M, Tomaszewski KJ, Bonus T, et al. Asthma that is not well-controlled is associated with increased healthcare utilization and decreased quality of life. J Asthma Off J Assoc Care Asthma. 2011 Mar;48(2):126–32.

15. Stanford RH, Gilsenan AW, Ziemiecki R, Zhou X, Lincourt WR, Ortega H. Predictors of uncontrolled asthma in adult and pediatric patients: analysis of the Asthma Control Characteristics and Prevalence Survey Studies (ACCESS). J Asthma Off J Assoc Care Asthma. 2010 Apr;47(3):257–62.

16. Nguyen K, Zahran H, Iqbal S, Peng J, Boulay E. Factors associated with asthma control among adults in five New England states, 2006-2007. J Asthma Off J Assoc Care Asthma. 2011 Aug;48(6):581–8.

17. Guilbert TW, Garris C, Jhingran P, Bonafede M, Tomaszewski KJ, Bonus T, et al. Asthma that is not well-controlled is associated with increased healthcare utilization and decreased quality of life. J Asthma. 2011;48(2):126–132.

18. Vietri J, Burslem K, Su J. Poor Asthma control among US workers: Health-related quality of life, work impairment, and health care use. J Occup Environ Med. 2014;56(4):425–430.

19. Zahran HS, Bailey CM, Qin X, Moorman JE. Assessing asthma control and associated risk factors among persons with current asthma - findings from the child and adult Asthma Call-back Survey. J Asthma Off J Assoc Care Asthma. 2015 Apr;52(3):318–26.

20. Mosnaim G, Lee LK, Carpinella C, Ariely R, Gabriel S, Lugogo NL. The impact of uncontrolled asthma on quality of life among treated, adherent patients with persistent asthma. J Allergy Clin Immunol. 2018 Feb 1;141(2):AB222.

21. Sadatsafavi M, McTaggart-Cowan H, Chen W, Mark FitzGerald J, Economic Burden of Asthma (EBA) Study Group. Quality of Life and Asthma Symptom Control: Room for Improvement in Care and Measurement. Value Health J Int Soc Pharmacoeconomics Outcomes Res. 2015 Dec;18(8):1043–9.

22. Wang P, Fu AZ, Wee HL, Lee J, Tai ES, Thumboo J, et al. Predicting preference-based SF-6D index scores from the SF-8 health survey. Qual Life Res Int J Qual Life Asp Treat Care Rehabil. 2013 Sep;22(7):1675–83.

23. Nurmagambetov T, Kuwahara R, Garbe P. The Economic Burden of Asthma in the United States, 2008-2013. Ann Am Thorac Soc. 2018 Mar;15(3):348–56.

24. Nurmagambetov T, Khavjou O, Murphy L, Orenstein D. State-level medical and absenteeism cost of asthma in the United States. J Asthma Off J Assoc Care Asthma. 2017 May;54(4):357–70.

25. Statistics UB of L. Highlights of Women's Earnings in 2012. 2002;

26. Nathan RA, Sorkness CA, Kosinski M, Schatz M, Li JT, Marcus P, et al. Development of the asthma control test: a survey for assessing asthma control. J Allergy Clin Immunol. 2004 Jan;113(1):59–65.
